# Evolutionary principles underlying neuron subtype encoding and diversification in animals

**DOI:** 10.64898/2026.04.13.718258

**Authors:** J. Hehmeyer, D. Harris, C.J. Lowe, H. Marlow

**Affiliations:** Department of Organismal Biology and Anatomy, University of Chicago, Chicago, IL, USA; Integrative Biology Program, University of Chicago, Chicago, IL, USA; Department of Biology, Hopkins Marine Station, Stanford University, Pacific Grove, CA, USA

## Abstract

Animals harbor a diversity of terminally differentiated cell types. This cell type diversity is particularly spectacular in the nervous system. While the shared evolutionary origin of the bilaterian neuron has been increasingly supported, it remains poorly understood when the first neuron diversified into the many types present in extant lineages and what molecular mechanisms accompanied this neuron diversification. Existing models of neuronal evolution are based on select observations from flies, nematodes, and humans, rather than systems-level comparisons across whole nervous systems and diverse species. Applying single-cell RNA-sequencing in the acorn worm *Saccoglossus kowalevskii* and performing cell type comparisons across protostome and deuterostome species, we show that neuronal subtypes in one species bear little resemblance to those found in different phyla. Previously identified cross-phyletic transcriptional similarities represent exceptions to a global pattern of divergence. Specifically, we observe that all neurons share the expression of genes associated with general features such as the presynapse and axon cytoskeleton, but distantly related species have evolved distinct usage of subtype-defining genes: they co-express unique combinations of neurotransmitter pathways, ion channels, postsynaptic receptors, and transcription factors. Additionally, we recover signal for an ancient neuronal regulatory code characterized by a strong enrichment for homeodomain transcription factors. Overall, these results suggest that while neurons largely share a core set of transcriptional features, a great deal of reshuffling of gene expression has occurred, supporting large-scale turnover in neuron subtype identities amongst major animal lineages.

## Introduction

The animal nervous system is one of the most conspicuous examples of how evolutionary processes have resulted in the emergence of a complex system. Vertebrates, arthropods, molluscs, and members of many other animal lineages in the superphylum Bilateria demonstrate highly complex nervous systems, consisting of dozens to thousands of morphologically and molecularly distinct neuron cell types (Z. Yao et al. 2023; Fincher et al. 2018; Taylor et al. 2021; Davie et al. 2018; Verasztó et al. 2025; Gavriouchkina et al. 2025; Schlegel et al. 2024) organized in networks of connectivity within diverse neural tissues (Figure 1A) (Schmidt-Rhaesa et al. 2015; Hejnol and Lowe 2015; Butler and Hodos 2005). Bilaterian nervous systems are united by many shared anatomical and functional properties, such as the somatic-axonic structure of neurons and the use of electrical and chemical signaling mechanisms to rapidly regulate the activity of neural and non-neural cells. At the same time, nervous systems are also highly structurally and functionally specialized: each species’ nervous system is closely integrated into the overall organization of its body plan and highly adapted to its unique environmental context and behavioral requirements. Thus, reconstructing when and how various features of nervous systems originated and changed promises to provide insight into both universal and unique mechanisms underlying the formation and function of these complex systems.

**Figure 1:**
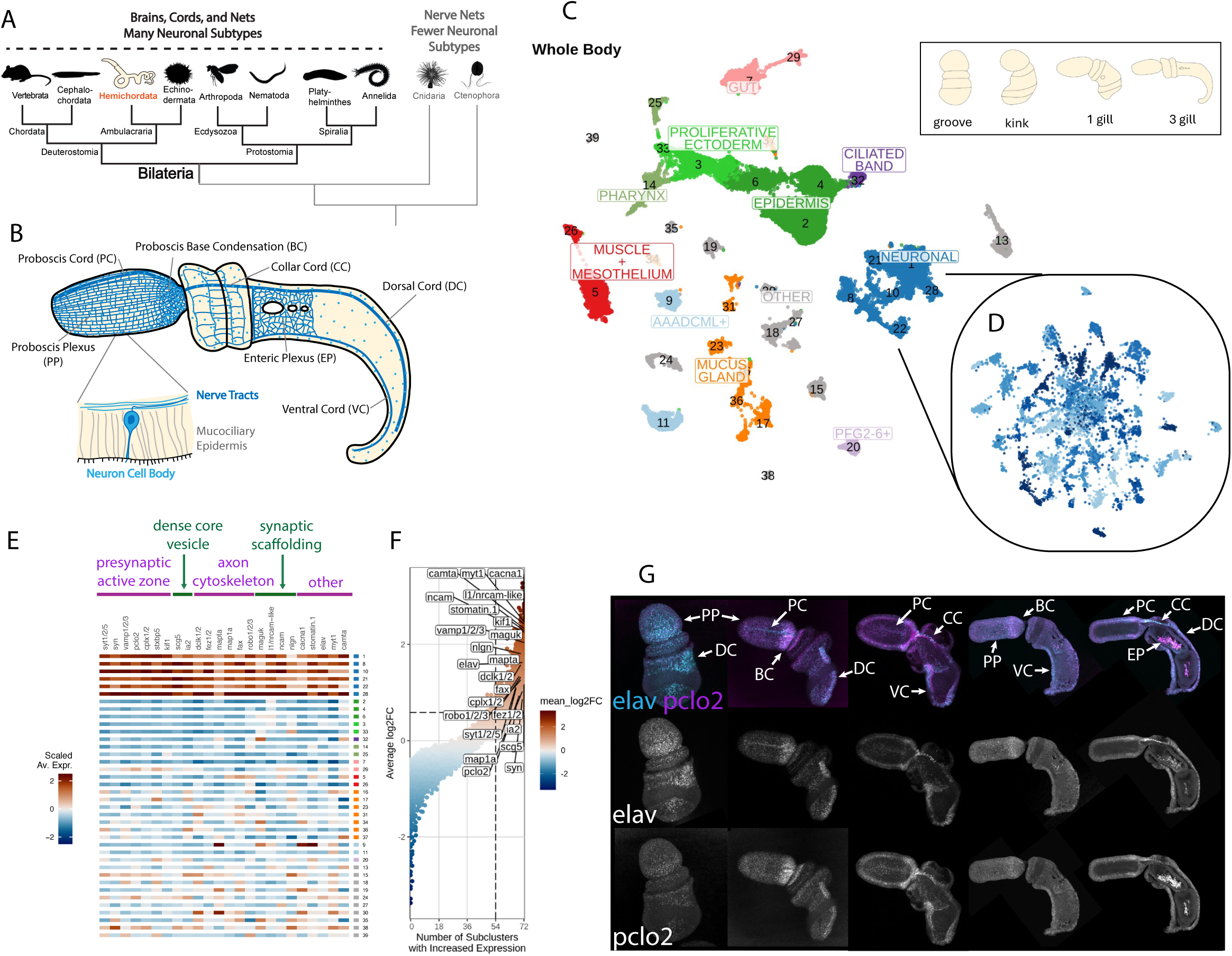
Whole-body single cell RNA-seq atlas of *Saccoglossus kowalevskii* enables the identification of neurons organism-wide and the identification of a pan-neuronal gene expression program. A). Phylogenetic tree of Metazoa highlighting the position of Hemicordata (beige) within Bilateria. Ambulacraria, including hemichordates, is sister to the chordates, which include cephalochordates and vertebrates. B). 3 gill-slit juvenile of *S. kowalevskii*. Three major body regions, proboscis, collar and trunk, make up the body plan and neurons are found in all three regions. Notable neural features are indicated: proboscis plexus (PP), proboscis cord (PC), proboscis base condensation (BC), collar cord (CC), dorsal cord (DC), ventral cord (VC), and enteric plexus (EP). C). UMAP of a whole-body single-cell atlas consisting of the stages illustrated (inset). Projections of individual stages onto the total clustering are presented in **Supplemental Figure 1**. Neurons represent a set of clusters indicated in blue. The late embryonic and juvenile stages of *S. kowalevskii* that are represented in the data include the groove, kink, 1-gill pair juvenile and 3-gill pair juvenile (inset) D). A re-clustering of all neurons produces 72 distinct neuronal clusters. E) Heatmap of neuronal markers detected across total cell clustering. Neuron clusters are indicated by blue squares accompanying cluster number on right side of panel. F). Plot showing the proportion of neuronal sub-clusters (x-axis) from the neuron-specific clustering with the average log-fold enrichment of identified neuronal markers indicated (y-axis). Each point represents a different gene. G). Hybridization chain reaction (HCR) *in situs* of RNA probes in groove embryos and 1-gill and 3-gill *S. kowalevskii* juveniles. Embryos and juveniles are double-stained with probes against *elav* and *picolo2* RNAs. At groove stage, expression is detected in the early proboscis plexus and dorsal cord, with additional expression domains detected by the 1-gill stage (base condensation, collar cord, proboscis cord, ventral cord) or 3-gill stage (enteric plexus) as the nervous system continues to develop.

While much emphasis has been placed on the structure, connectivity and function of the many different formats of nervous systems, molecular comparisons have emerged as powerful tools in understanding the evolutionary history of its fundamental units, neurons. The homology of many neural features has been established by studies integrating genomic and functional data from across the animal tree. Based on similarities in the presence, expression, and function of genes, these studies have revealed the evolutionary homology of many core cellular features of bilaterian neurons, such as synaptic vesicles, the presynaptic active zone, and postsynaptic scaffolding and receptor networks (Burkhardt and Jékely 2021; T. J. Ryan and Grant 2009; Burkhardt and Sprecher 2017; Arendt 2020; Rosenthal et al. 2025). These studies demonstrate that many of the components needed to build a neuron, generally, are ancient and conserved. In contrast, it is still unknown when and how in evolutionary history the plethora of molecularly distinct neurons observable in any given species arose.

Candidate gene-based approaches have identified cases of gene co-expression conserved across distantly related species from separate phyla, suggesting that there could be cross-phyletic conservation of neuron identity (Nomaksteinsky et al. 2013; Lloret-Fernández et al. 2018; Flames and Hobert 2009; Remesal et al. 2020; Catela et al. 2019; Popsuj and Stolfi 2021; Denes et al. 2007; Nakamura et al. 2023; Arendt et al. 2021; Tessmar-Raible et al. 2007). Accordingly, it has been suggested that many neuron populations and the gene expression defining their core identities can be traced back to the bilaterian last common ancestor (Arendt et al. 2016, 2019). In contrast, recent comparisons at the level of full transcriptomes have revealed that most neuron subtypes lack any cross-phylum counterpart that is any more similar than any other neuron (Gavriouchkina et al. 2025; Styfhals et al. 2022; Dai et al. 2024); these results are more consistent with alternate evolutionary histories such as independent subtype expansions (Hehmeyer et al. 2024) or multiple origins of neurons from non-neural cell types (Moroz 2021).

The systems-level mechanisms of neuronal diversification are similarly poorly understood. Developmental and regulatory programs need to be structured to accommodate the generation of a tremendous diversity of subtypes. Some classes of transcription factors have been found to be expressed during neural development in diverse species (Hobert and Westphal 2000; Hobert 2025; Srivastava et al. 2010; C. Xu et al. 2024; Plessier and Marlow 2026; Richards and Rentzsch 2015), a pattern which may relate to the regulatory basis by which the genome encodes a multitude of subtype identities. For example, nematodes deploy a combinatorial code of homeodomain transcription factors that cooperatively distinguish neuron types by activating unique but overlapping sets of subtype-function-defining genes; homeodomains are often expressed in mature neurons in other bilaterian species, though whether they serve the same purpose is unknown (Hobert 2021). However, it remains unknown the extent to which common or unique regulatory mechanisms are utilized across diverse animal lineages to encode a diversity of neurons.

The phylum Hemichordata is a critical taxon for reconstructing the evolutionary history of bilaterian nervous systems. Due to its phylogenetic position within Deuterostomia (**Figure 1A**), it serves as a phylogenetic bridge between the well-characterized nervous systems of vertebrates and model protostomes such as insects and nematodes, allowing for a more informed reconstruction of the evolution of neurons across all lineages with complex nervous systems. Here, we leverage single-cell RNA sequencing (scRNA-seq) to generate a whole body single-cell atlas for *Saccoglossus kowalevskii*, a species of hemichordate worm. We set out to identify the gene expression principles underlying neuronal identity and subtype diversity in this species, then assess whether global comparisons of neurons across a diverse sampling of bilaterian species would allow us to identify a common set of neuronal features largely shared among lineages, and whether a granular reconstruction could be achieved in which homologous subpopulations of neurons might be identified within Bilateria. We observe strong cross-species similarities in the sets of genes that unite neurons as a class and distinguish individual subtypes from each other. We also identify a common regulatory grammar of transcription factors (TFs) that is deployed across neurons that relies heavily on homeodomain-family genes. Yet, we find no evidence to strongly support homology among subtypes of neurons between phyla; instead, our analysis suggests that the evolution of neurons involved pervasive changes in the patterns of usage of an old neuronal gene toolkit, resulting in a poor evolutionary maintenance of any ancient subtype identities as well as contributing to lineage-specific neural diversifications.

## Results

### Identification of hemichordate neurons and pan-neuronal features

Previous morphological studies and targeted molecular characterization of *S. kowalevskii* neurons has provided a first snapshot of the organization of hemichordate nervous systems (Andrade López et al. 2023), but the comprehensive characterization of neural types needed to perform comparative analyses with other lineages has thus far been lacking. In order to systemically identify and characterize the gene expression profiles of *S. kowalevskii* neurons in an unbiased manner, we implemented scRNA-seq. To capture cells from throughout the diffuse and broadly distributed nervous system of this animal (**Figure 1B**), as well as to allow for comparison of neurons to other cell types, we elected to generate our atlas using whole body samples, specifically working with individuals in the late embryonic and juvenile stages of development (**Figure 1C**, upper right), the stages spanning neuron terminal differentiation in this species (Cunningham and Casey 2014; Kaul and Stach 2010; Andrade López et al. 2023). We generated new data from juvenile hemichordates using the 10X Chromium droplet-based platform and re-analyzed existing embryonic and juvenile data (Bastide et al. in review) generated using the plate-based MARS-seq approach. Data were mapped to a corrected gene model set (outlined in methods) and then quality filtered using a multistage preprocessing pipeline that includes steps to detect and remove anucleated debris from the droplet-based datasets (Montserrat-Ayuso and Esteve-Codina 2024; Muskovic and Powell 2021) (**Supplemental Figure 1A**). Following cross-sample integration and clustering of our data, we recovered ∼37,000 high-quality cells comprising 39 clusters (**Figure 1C**, center), with each cluster represented by cells from multiple stages of development and from both single-cell platforms (**Supplemental Figure 1B-C**), confirming that integration did not suffer from method-dependent batch effect.

To begin to characterize neuronal gene expression programs in hemichordate neurons, we first needed to distinguish neurons from other cells. As a first step, we annotated clusters based on the expression of previously characterized genes from *Saccoglossus kowalevskii* and other hemichordates (Simakov et al. 2015; Cunningham and Casey 2014; Christopher J. Lowe et al. 2003; Fritzenwanker et al. 2019; C. J. Lowe et al. 2006; Sébastien Darras et al. 2018; Pani et al. 2012; Fritzenwanker et al. 2014; S. Darras et al. 2011; Satoh et al. 2014; Tagawa et al. 2014; Ikuta et al. 2013; Vianello et al. 2026; Andrade López et al. 2023; Gillis et al. 2011; C. Chou et al. 2024; Gąsiorowski et al. 2021; Su et al. 2019; Green et al. 2013; Okai et al. 2000; Aronowicz and Lowe 2006) (**Supplemental Tables 1-2**). We were able to classify the majority of cell clusters into several major classes: neurons, proliferative ectoderm, differentiated epidermis, ciliated band, pharynx (cells from the mouth, stomochord, and gill tissues), gut (cells from esophageal, intestinal, and hindgut tissues, including digestive cells and hepatic-like cells), muscle and mesothelium (including podocytes), and mucus gland cells (**Figure 1C**, **Supplemental Figure 2**). We also identified clusters corresponding to previously characterized hemichordate cell populations with known anatomical distribution but less well understood function. These are the AAADCmL+ cells, known to be scattered throughout the epidermis (Simakov et al. 2015), which share some features with arthropod glia (*gcm*+, *glul*+, *ebony+*) (Jones et al. 1995; Hosoya et al. 1995; Freeman et al. 2003; Pantalia et al. 2023); and the cells embedded in the pharyngeal epithelium that express the *PfG2/3/4/6*-type lectin genes (Okai et al. 2000) (**Figure 1C**, **Supplemental Figure 2**).

As the annotation of a cell type based only on a handful of previously identified markers risks the erroneous inclusion of irrelevant cells or the accidental exclusion of relevant populations of interest, we implemented a series of analyses to carefully and systematically identify neurons. Previous work on *Saccoglossus* has identified several genes that show broad and specific expression in neural tissues in hemichordates, including the presynaptic vesicle complex components synapsin (*syn*) and synaptotagmin 1/2/5 (*syt1/2/5*); doublecortin-like kinase (*dclk1/2*), a conserved regulator of axon microtubule structure; and the RNA-binding protein gene *elav* (Cunningham and Casey 2014; Andrade López et al. 2023; Christopher J. Lowe et al. 2003; Nomaksteinsky et al. 2013). We identified 6 clusters in the full cell atlas that show high expression of these neuronal markers (**Figure 1E**), our putative neuron clusters. As a first step toward validating this classification, we carried out differential expression analysis to identify genes with increased expression in each cluster relative to other cells. We identified 50+ significantly positively differentially expressed genes (“marker genes”) per cluster (**Supplemental Table 3**). The putative neuronal clusters show increased expression of dozens of genes with well-characterized core neuronal functions in other species (**Figure 1E**, **Supplemental Table 3**). Specifically, marker genes shared amongst most or all of our putative neuron clusters include components of synaptic vesicles and the cytomatrix at the presynaptic active zone (*syt1/2/5, vamp1/2/3 (synaptobrevin), pclo2, snap23/25, syde1/2, syn, syt4/11, cadps (caps1), cplx1/2*, *stxbp5* (*tomosyn*)) (Burkhardt and Sprecher 2017; Wentzel et al. 2013; Wolfes and Dean 2020; Gundelfinger et al. 2016; Fujita et al. 1998); genes associated with dense-core vesicles (*scg5 (7b2), ia2 (ptprn)*) (Cai et al. 2011; Harashima et al. 2005; Stojilkovic et al. 2025; MBIKAY et al. 2001); components or regulators of the axon cytoskeleton (*kif1, kif5, fez1/2 (unc-76), mapt (tau)*, the map1a/b/s paralog *map1a, dclk1/2*, the faxc paralog *fax1*, *robo1/2/3*) (Okada et al. 1995; Campbell et al. 2014; Bloom and Horvitz 1997; Gindhart et al. 2003; Nawabi et al. 2015; Hill et al. 1995; Gonda et al. 2020; Cario and Berger 2023); genes involved in scaffolding the synapse at the pre- or post-synaptic membrane (*maguk, nrcam/l1cam-like, ncam, nlgn*) (Giagtzoglou et al. 2009; Liebeskind et al. 2017; Zhu et al. 2016; Sakurai 2012); the voltage-gated calcium channel *cacna1* (Szymanowicz et al. 2024); the ion channel interacting gene *stomatin* (*stom*) (Lapatsina et al. 2012); and several TFs, including *myt1/2/3* (orthologous to vertebrate *myt1*, *myt1l* (*myt2*), and *st18* (*myt3*)), which has a conserved role in repressing non-neuronal gene programs (J. Lee et al. 2019; Mall et al. 2017), and *nr4a* and *camta*, unrelated TFs which all play roles in activity-dependent transcriptional regulation (Bas-Orth et al. 2016; Hawk and Abel 2011), confirming that these cells do indeed have a neuronal identity. Consistent with these observations, the set of genes with enriched expression in the putative neuron clusters demonstrate high representation of neuronally-associated GO terms such as "synapse", "axon", "monoatomic ion-gated channel activity", "modulation of chemical synaptic transmission", and "neurotransmitter transport" (**Supplemental Figure 3A**, **Supplemental Table 4**). Critically, no other clusters demonstrate high enrichment for all of these GO terms (**Supplemental Figure 3A**). Thus, only our neuronal clusters show transcriptional signatures clearly indicative of neuronal function.

In addition to this expanded marker approach, we utilized evolutionary comparisons to more broadly assess cell types for inclusion in our neuronal cell set. SAMap integrates scRNA-seq across species, allowing for the identification of clusters from different species that express similar sets of genes (Tarashansky et al. 2021). We used SAMap to compare our *Saccoglossus* cells to those from comprehensive, well-annotated whole-body single cell atlases from zebrafish (*Danio rerio*) (Sur et al. 2023) and fly (*Drosophila melanogaster*) (H. Li et al. 2022). As a proof of principle, we were able to recover known, ancient class-level relationships amongst certain populations (muscle, gut, proliferative populations) (Jingjing Wang et al. 2021; Tarashansky et al. 2021; Piovani et al. 2023; Robertson et al. 2024) (**Supplemental Figures 3B**, **Supplemental Table 5**). Importantly, *Saccogossus* neuron clusters annotated using the expanded marker approach also demonstrated high similarity to neurons from the other species, whereas *Saccoglossus* clusters that are not annotated as neurons showed no such similarity (**Supplemental Figure 3B**, **Supplemental Table 5**). Altogether, these approaches allowed us to confidently conclude that we had included all neuron-like cells for subsequent analyses.

Beyond allowing us to distinguish neurons from non-neuronal cells, these analyses suggested the existence of a core set of global features shared across *Saccoglossus* neurons. To break down the neuron clusters into their underlying subpopulations so that we would be able to identify the features uniting and distinguishing each of them, we carried out subclustering of the neuron clusters, recovering 72 subtypes of 40 to 300 cells each (**Figure 1D**). We observed that many of the neuron marker genes we identified—including presynaptic complex genes, axon cytoskeleton genes, and synaptic scaffolding genes—show increased expression across a majority of neuronal subclusters relative to their “background” expression level in non-neuronal cells (**Figure 1F**). Thus, we are confidently able to identify a core set of molecular features that define a hemichordate neuron.

Having identified neurons in the scRNA-seq data, we next set out to validate that these cell types correspond to morphologically identifiable neural populations in tissue and that these markers are indeed co-expressed in cells. We utilized HCR *in situ* hybridization, confirming that the newly identified hemichordate broad neuronal marker *pclo2* is co-expressed with *elav*, though with different subcellular localization, throughout embryonic and juvenile neural tissues (**Figure 1G**). Overall, the expression of the two genes closely matches the known distribution of neuronal cells in this animal over ontology (Cunningham and Casey 2014; Andrade López et al. 2023; Kaul-Strehlow and Stach 2013). Thus, we are able to confidently identify neuronal cells in our hemichordate cell atlas which are consistent with the known organization and location of neurons in juvenile hemichordate worms. This provides a powerful entry point to further characterizing transcriptional features of these neurons that relate to their development and evolution.

### Molecular Complexity of the Hemichordae Nerve Net

We next set out to characterize the transcriptional diversity among our 72 subclusters of hemichordate neurons (**Figure 2A**). We first assessed the patterns of usage of effector (non-TF) genes, those directly responsible for building and maintaining the functional identity of the cell. These genes contribute to the many axes/dimensions of neuronal function: wiring, electrical properties, chemical inputs and outputs, and other aspects of the phenotype. Traditionally, the vast functional diversity of neurons in the nervous systems of bilaterians such as vertebrates, insects, and nematodes is practically simplified by classifying each neuron by the neurotransmitter(s) it releases at the synapse. However, recent studies have suggested that differences in the presence and usage (i.e. small molecule vs. peptide, single molecule vs. co-transmission) of neurotransmitter systems exist between animal lineages (Goulty et al. 2023; Bauknecht and Jékely 2017; Hardege et al. 2022; Burkhardt and Jékely 2021). Thus, important questions have remained about when each of these transmitter classes arose in animals and how they are utilized in disparate nervous systems .

**Figure 2:**
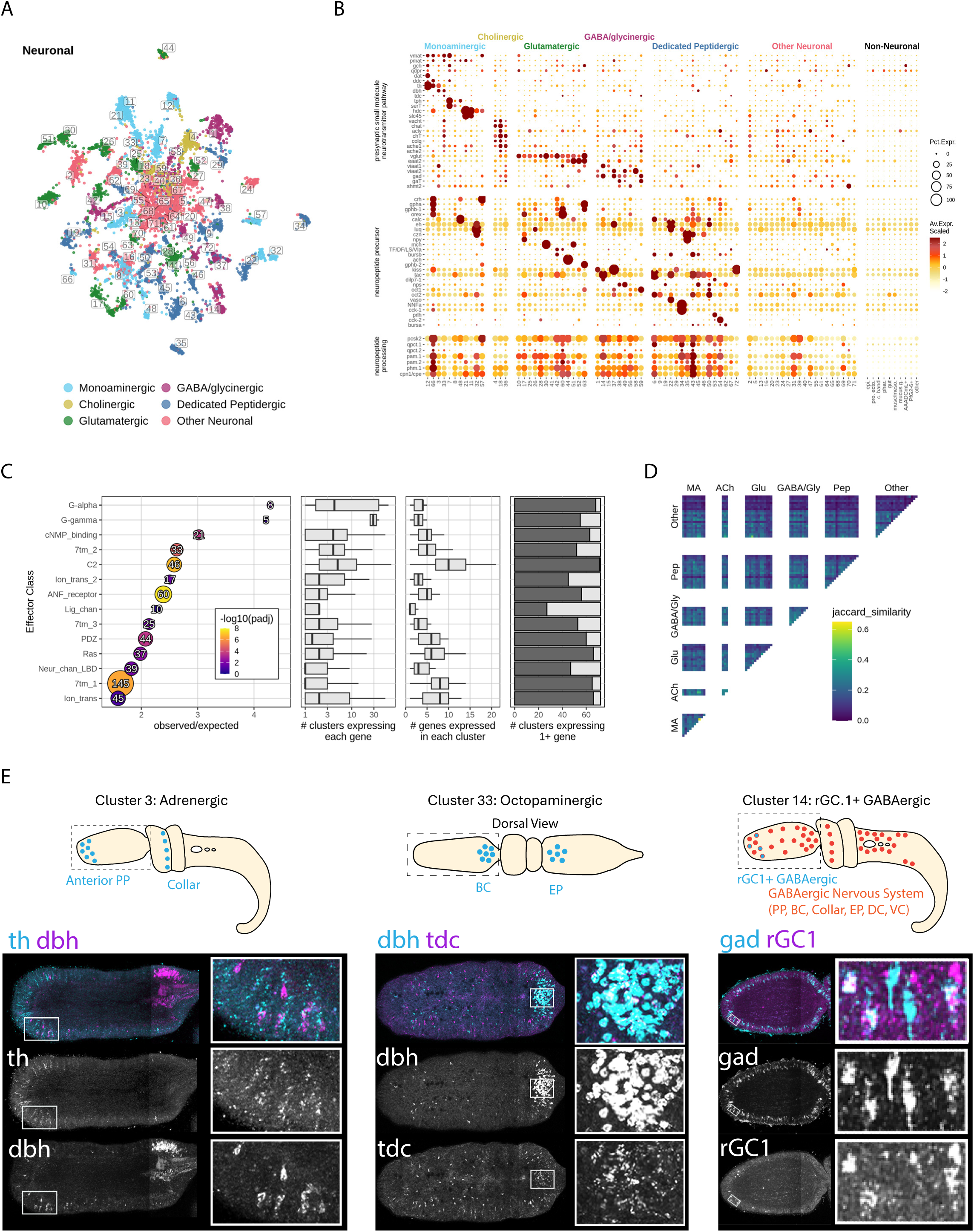
Neuron-specific clustering analysis identifies a high transcriptional diversity of sub-classes of hemichordate neurons. A). UMAP projection of neuron-specific clustering color-coded by neurotransmitter class. B). Dot plot showing expression of markers informative for neurotransmitter classes. C) Identification of gene families with enriched expression in neurons and characterization of their expression breadth. Left: Enrichment of Pfam domain-based gene families amongst neuronally expressed genes (genes with enriched expression in at least one neuron subcluster); significance of enrichment and number of neuronally expressed genes indicated by size and color of points. Middle: Bar plot showing the number of neuron subclusters that express at least one gene from each Pfam class. Right: Box plot showing the number of clusters expressing each gene. D). Similarity between pairs of neuron subclusters based on enriched expressed genes. E). Schematics of the anatomical locations of three individual neuron subclusters (cluster 3: adrenergic neurons; cluster 33: octopaminergic neurons; and cluster 14: rGC1+ GABAergic neurons) in the *S. kowalevskii* juvenile and corresponding *in situ* RNA-HCR images in 3-gill stage worms for probe combinations specific to each subcluster. Left: probes for *dopamine beta-hydroxylase* (*dbh*, cyan) and *tyrosine hydroxylase* (th, magenta) in the juvenile. Within the proboscis, *dbh* expression overlaps with *th* expression only in the anterior proboscis plexus (white box and inset). Middle: probes for *dopamine beta-hydroxylase* (*dbh*, magenta) and *tyrosine decarboxylase* (*tdc*, magenta) in the proboscis. Within the proboscis, most cells co-expressing dbh and *tdc* expression are found in the region of dorsal proboscis base condensation (white box and inset). Right: probes for *receptor guanylate cyclase 1* (*rGC1*, magenta) and *glutamate decarboxylase* (*gad*, cyan) in the proboscis. *rGC*+, *gad*+ cells are restricted to the anterior proboscis (white box and inset).

Thus, we elected to assess the expression of known neurotransmitter-associated genes, both to use this as a starting point to understand hemichordate neuronal diversity, and to gain further insight into the evolutionary history of neurotransmitter systems. The existence of numerous distinct small molecule and peptidergic neuron populations in hemichordates was previously investigated *in situ* by imaging work using histochemical staining, cross-reactive antibodies, and single- or double-gene RNA labeling (Andrade López et al. 2023; Nomaksteinsky et al. 2009; Dautov and Nezlin 1992; Cunningham and Casey 2014; Kaul-Strehlow et al. 2015), but these studies did not test for the presence of all known small molecules or neuropeptides, comprehensively assess co-transmission, nor resolve subtype diversity within these neurotransmitter classes. We manually curated and examined the expression of genes diagnostic of presynaptic neurotransmitter usage, that is, the genes responsible for the synthesis, secretion (vesicular loading), and cytoplasmic reuptake of small molecule neurotransmitters, as well as those responsible for the production of peptide neurotransmitters (Miyajima et al. 2022; Ichinose et al. 2008; Hwang et al. 1998; Zeisel et al. 2018; Goulty et al. 2023; C. Wang et al. 2024; Puttonen et al. 2017; Y. Xu and Wang 2019; Duan and Wang 2010; Pörzgen et al. 2001; Zafra et al. 1997; Beigneux et al. 2004; Ito et al. 2012; Wei et al. 2003; Zhang et al. 2010; Busby et al. 1987; Fischer and Spiess 1987; Han et al. 2004) (see **Supplemental Table 6** for full annotation of functions of pathway genes). Based on expression of these genes, we classified hemichordate neurons into 1 of 4 small molecule neurotransmitter classes: monoaminergic, cholinergic, glutamatergic, and GABA/glycinergic (**Figure 2A-B**, **Supplemental Table 6**). Of particular note, we observe evidence for the release of a diversity of distinct monoamine molecules, with the hemichordate monoaminergic system showing some features once believed to be restricted to protostomes. Distinct dopamine, noradrenaline, octopamine, serotonin, and histamine -releasing populations can be distinguished by their differential, combinatorial expression of monoaminergic pathway genes (**Supplemental Table 6**). Our detection of separate adrenergic and octopaminergic neurons, once believed to represent functionally analogous populations in vertebrates and protostomes (Pflüger and Stevenson 2005; Verlinden et al. 2010; Roeder 1999) is consistent with the previous functional characterization of unique receptors for these molecules in the *Saccoglossus* genome, and adds to the growing recognition of the coexistence of these systems in invertebrate bilaterians (Ruppert et al. 2024; Bauknecht and Jékely 2017; Jin et al. 2023). Moreover, histamine vesicular loading is carried out in some hemichordate histaminergic populations by a solute carrier family 45 gene (*slc45-like*), as in arthropods (Y. Xu and Wang 2019), and in other populations by vesicular monoamine transporter (*vmat)*, as occurs in vertebrates (Puttonen et al. 2017). Altogether, this data indicates that the origin of many of the monoamine processing and transport genes along the bilaterian stem (Goulty et al. 2023; Burkhardt and Jékely 2021) was followed by their canonical usage in at least five distinct pathways prior to the divergence of extant bilaterian lineages, and there has generally been conservation of this ancestral monoaminergic system in hemichordates, in contrast to other phyla which show differential losses of some ancient components.

In addition to small molecules, nervous systems also rely heavily on neuropeptides for fast or modulatory transmission (Lingueglia et al. 1995; Assmann et al. 2014; Schmidt et al. 2018; Ripoll-Sánchez et al. 2023; Jékely 2021; Ceballos et al. 2024; Plessier and Marlow 2026). We find that the genes responsible for neuropeptide precursor processing generally show high expression across all neuron populations (**Figure 2C**, middle). In contrast, analyzing the expression of previously identified peptide precursor genes (**Supplemental Table 7**), we find that many *Saccoglossus* neurons release one, two, or even several neuropeptides, but with each neuropeptide expressed in a relatively small number of distinct populations (**Figure 2B**). While all hemichordate neurons express the machinery required for neuropeptide synthesis, not all are found to express annotated neuropeptide genes, a discrepancy likely due the incomplete annotation of the neuropeptidome due to the lack of peptidomics studies in hemichordates. Nonetheless, our observations reveal a complex, combinatorial expression of a diversity of peptides in the hemichordate nervous system.

Though we detect multiple peptidergic clusters that lack any detectable expression of presynaptic small molecule neurotransmitter genes, populations we refer to here as “dedicated peptidergic” (**Figure 2A-B**), we find that the majority of neuropeptides are, in fact, highly expressed in some glutamatergic, GABA/glycinergic, and monoaminergic populations. In contrast to our observations of peptide-peptide and peptide-small molecule co-transmission, we detect very few cases of strong co-expression of genes implicated in the presynaptic usage of different small molecules, an observation distinct from the diverse instances of co-release of small molecules in other bilaterians (Z. Yao et al. 2023; Tritsch et al. 2016; C. Wang et al. 2024; Wallace and Sabatini 2023). Altogether, these results provide evidence that dual small molecule-peptide transmission is the dominant transmitter phenotype of hemichordate neurons.

While neurotransmitter usage is the conventional axis along which to classify neuron types, work in other nervous systems indicates that there is significant heterogeneity within neurotransmitter classes, such that neurons with similar neurotransmitter usage can be just as molecularly distinct from each other as they are from neurons expressing different neurotransmitters (see, for example: (Zeisel et al. 2018; Z. Yao et al. 2023; Kratsios and Hobert 2018; Konstantinides et al. 2018)). We carried out differential expression analysis to identify genes with significantly increased expression in each neuronal subcluster relative to their expression in non-neuronal cells (**Supplemental Table 8**). Our differential expression analysis demonstrates that multiple other aspects of neuron identity beyond neurotransmitter pathway genes distinguish the *Saccoglossus* neuron populations. To characterize the genes contributing to these differences at a systems level, we assessed the Pfam domains over-represented amongst genes with significantly enriched expression in one or more neuron subclusters (**Supplemental Table 9**). We found that highly over-represented domains include those with known roles in neuron-type-specific function in vertebrates, flies and nematodes (**Figure 2C**). These include the domains defining GPCRs and G-proteins (7tm_1, 7tm_2, 7tm_3, G-alpha, G-gamma, ANF_receptor) (Krishnan et al. 2013; X. Liu et al. 2004; Hille 1992), other metabotropic receptors (ANF_receptor) (Ortiz et al. 2006; Turek et al. 2023), ion channels (ion_trans, ion_trans_2) (Davila-Velderrain and van Giesen 2025), ionotropic receptors (ANF_receptor, Neur_chan_LBD, Lig_channel) (Kuryatov et al. 1994; Yang et al. 2025; Zhou et al. 2021; Apostolakou et al. 2022), and genes involved in postsynaptic and ion channel interaction and signaling networks (PDZ, Ras) (Zhu et al. 2016; Feng and Zhang 2009; Gu and Stornetta 2007; Wilkinson and Coba 2020). At least one gene with each of these domains can be detected in the majority of neuronal subclusters, but each individual gene’s expression is restricted to a limited number of subclusters (**Figure 2C**). Indeed, due to the differential and combinatorial patterns of expression of these effectors, neurons expressing the same neurotransmitter are not any more similar to each other than to neurons expressing different neurotransmitters–e.g. glutamatergic neurons are no more similar to one another than they are to GABAergic or monoaminergic neurons (**Figure 2D**).

Altogether, our results indicate that there is a high transcriptional diversity of neurons in hemichordate. Each of the subclusters identified here represents a highly distinct subpopulation (i.e. neuronal subtype), as can be visualized by our identification of the distribution of adrenergic neurons, octopaminergic neurons, and rGC+ GABAergic neurons within the juvenile nervous system (**Figure 2E**). Subtype identity is defined by multiple layers of effector genes, with these genes largely of the same functional classes as in flies, nematodes and vertebrates, despite the largely non-centralized nature of the acorn worm nervous system. Having characterized the effector diversity of hemichordate neurons, we next set out to understand the contribution of transcription factor expression to transcriptionally distinguishing this neuronal diversity.

### A homeodomain Transcription Factor code for hemichordate neurons

Cellular identity is established and maintained through the activity of unique combinations of transcription factors. To characterize the TF regulatory programs of hemichordate neuron populations system-wide, we identified and annotated hemichordate TF genes based on domain composition, and then classified these genes into 12 major families: C2H2-type zinc finger (zf-C2H2); homeodomain (HD); Forkhead; helix-loop-helix (HLH); bZIP; high mobility group, including Sox-like (HMG/Sox); Ets; GATA; Nuclear Receptor zinc finger (Nucl.Rec.); T-box; Myb; and "other", a designation for TFs not falling into these other major families (**Supplemental Table 10**). The number of TFs in each of these families is largely consistent with a previous characterization of the hemichordate TF complement (de Mendoza and Sebé-Pedrós 2019). Of the 1,168 TFs we predict in the genome, we detected the expression of 965 (82%) in at least 5 cells (**Supplemental Table 10**).

We assessed the expression of the *Saccoglossus* TFs across neuronal subclusters (**Figure 3A** and **Supplemental Table 11**). We find that most subclusters express 10 or more TF genes, largely from a mix of different families (**Figure 3A**, bottom). Moreover, the expression of the majority of TFs is restricted to just one or a few clusters, with a small number of TFs showing broader patterns of expression (**Figure 3B**). The TF programs of each cluster are entirely unique and can be confirmed *in situ*, for example, cells representing the *vasotocin*+ peptidergic cluster 22 uniquely express *hbn* (**Supplemental Figure 4A**), whereas octopaminergic neurons (cluster 33) uniquely express *fer2* (**Supplemental Figure 4B**).

**Figure 3:**
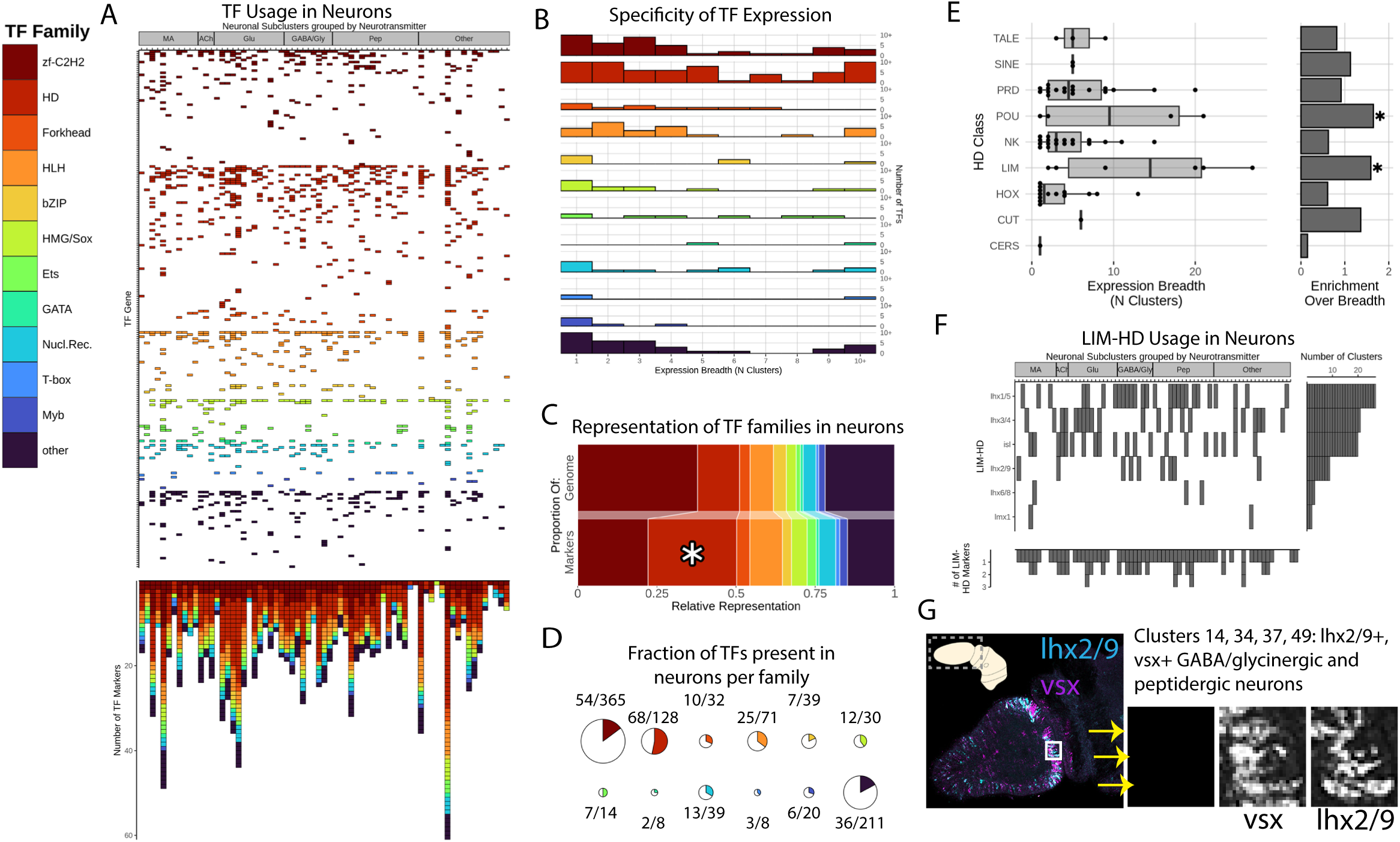
Homeodomains are enriched amongst neuronally-expressed TFs. A). Transcription factor expression across neuronal subtypes. Neuron subclusters (x-axis) are ordered by transmitter class. Transcription factors (y-axis) are ordered and colored by family (left). Stacked bar plots show the number of genes from each TF family which are detected in each subcluster. B). Bar plots show the number of TFs (y-axis) with each possible expression breadth value (x-axis). C). Relative representation of TFs contained in the genome (upper bar) versus those represented as markers of neurons (lower bar). Note the statistically significant over-representation of homeodomains as neuronal marker TFs. Asterisk indicates significant enrichment with adjusted p<0.05. D). Each colored fraction of a pie chart indicates the proportion of each TF family that is expressed in neurons. Total pie chart size is proportional to the size of the TF family. More than half of all homeodomain genes are detected in neurons. E). Differences in expression breadth amongst HD TFs of different classes. Left: Box plot showing the average number of neural cell clusters in which each class of HD TFs is expressed. Right: Enrichment for broad expression, normalized by the number of genes in each HD TF class for the data presented. Asterisks indicate significant enrichment with adjusted p<0.05. F). Broad and semi-complementary expression of LIM class HD TFs. Center: Expression of LIM class HD TFs (y-axis) in neuronal subclusters (x-axis), ordered by transmitter class. Right: The number of clusters each gene is detected in. Bottom: The number of LIM HD TFs expressed in each cell cluster. G). HCR *in situ* showing co-expression of transcripts of *lhx2/9* (cyan) and *vsx* (magenta) in the area of proboscis base condensation in a 1-gill stage *S. kowalevskii* juvenile (white box and insets).

The global neuronal TF complement shows evidence of similarity to that of other bilaterian species. This is apparent at the both level of individual genes, as well as at the more general level of TF families. The majority (53%) of the neuronal TFs have annotated neuronal functions in other species, as compared to 28% of TFs that are expressed only in non-neuronal cells (**Supplemental Figure 4C**). The HD TF family is significantly over-represented amongst neuronally expressed TFs (**Figure 3C**), an observation that has also been made in flies and nematodes (Özel et al. 2022; Hobert 2021). While many zf-C2H2 genes are also expressed, fewer than a third of those present in the genome are expressed in neurons, a pattern that holds for all other TF families with the exception of HDs (**Figure 3D**). HDs are also distinct in their discriminatory power: the expression of HDs alone is nearly sufficient to distinguish all subclusters, as 64 unique combinations of HDs can be detected amongst the 72 subclusters (**Supplemental Table 12**).

HD TFs include genes with both broad and restricted neuronal expression (**Figure 3B**). Members of the LIM and POU classes of HDs tend to show expression across many neuron subclusters, in contrast to the more restricted expression displayed by other HD genes (**Figure 3E**). Six of eight hemichordate LIM-HDs are expressed in neurons (**Supplemental Table 10**), and they show highly complementary patterns of broad expression: most of these genes are expressed in eight or more subclusters (**Figure 3E**, **Figure 3F**, right), and most subclusters express 1 or 2 LIM-HDs (**Figure 3F**, bottom), such that LIM-HD expression altogether encompasses almost all neuronal populations (**Figure 3F**). Consistently, *ldb1/2* (also known as *CLim*), a transcriptional cofactor which allows for the homo- and hetero-dimerization of LIM-HDs (G. Liu and Dean 2019), is enriched in many of the neuronal subclusters (**Supplemental Figure 5A**). Most subclusters can be distinguished by the combinatorial expression of LIM-HDs and 1-2 other HD TFs (**Supplemental Table 12**). By performing HCR experiments, we can identify the subset of the neurons co-expressing the LIM-HD *lhx2/9* and the HD *vsx* (**Figure 3G**, **Supplemental Figure 5B**). *lhx2/9*+ neurons can broken down in *vsx*+ and *vsx*- populations (roughly corresponding to GABA/glycinergic populations and non-GABA/glycinergic populations, respectively), with additional HDs distinguishing *lhx2/9*+,*vsx+ and lhx2/9*+,*vsx-* subtypes (**Supplemental Figure 5B**). Some of the LIM-HDs show rough associations with certain neurotransmitters, for example, *lhx1/5* and *lhx2/9* are more frequently associated with GABA/glycinergic neurons than with glutamatergic neurons, whereas the opposite is true for *lhx3/4* and *isl* (**Figure 3F**). However, overall, neurons of the same neurotransmitter type do not display more similar TF complements (**Supplemental Figure 5C**), in line with similar observations in other species (Closser et al. 2022; Kratsios and Hobert 2018; Konstantinides et al. 2018). Altogether, our analysis reveals that each hemichordate neuron cluster expresses a unique combination of TFs, driven largely by the combinatorial specificity of broadly expressed LIM HDs in conjunction with the more restricted expression of additional HD TF genes.

### Defining “neurons” across phyla

Previous work has established that neurons share a common origin in Bilateria, and most likely with Cnidaria, on the basis of global transcriptional similarity (**Supplemental Figure 3B-C**) (Sebé-Pedrós et al. 2018; Jingjing Wang et al. 2021; Tarashansky et al. 2021). We sought to more specifically pinpoint features of the Bilaterian neuronal signature and assess whether there is evidence for homology of specific subclasses of neurons by comparing hemichordate neurons to publicly available scRNA-seq datasets from 7 other bilaterian animals: the echinoderm *Paracentrotus lividus* (sea urchin) (Paganos et al. 2025); the chordates *Mus musculus* (mouse) (Zeisel et al. 2018), and *Branchiostoma floridae* (amphioxus) (Dai et al. 2024); the arthropod *Drosophila melanogaster* (fly) (D. Lee et al. 2025; Davie et al. 2018; Allen et al. 2020; Özel et al. 2021; Hopkins et al. 2023; H. Li et al. 2022); the nematode *Caenorhabditis elegans* (roundworm) (Taylor et al. 2021); the platyhelminth *Schmidtea mediterranea* (flatworm) (Fincher et al. 2018); and the annelid worm *Platynereis dumerilii* (ragworm) (Stockinger et al. 2024; Milivojev et al. 2025) (see Methods). We first carried out differential expression analysis as we did in *Saccoglossus* (each neuronal subcluster vs. non-neuronal “background”) (**Supplemental Table 13-14**); in parallel, we generated gene orthology assignments for these species (see Methods and **Supplemental Table 15**).

We then assessed whether we could identify the core set of neuronal effectors driving this cross-phyletic similarity in Bilateria as well as features which might unite more specific subtypes of neurons across phyla. We used a phylogenetic comparative approach (Methods) to identify orthogroups that tend to conserve broad neuronal expression across species (**Supplemental Table 16**). We found that the species in our analysis share the broad neuronal expression of genes with general roles in the pre-synapse, the axon, G-protein signaling, and neuropeptide processing and secretion (**Figure 4A**). Like we did with *Saccoglossus*, we identified the gene sets over-represented amongst neuronally expressed genes for each species. Again, we uncover a conserved pattern: across species, neuronally expressed genes show an enrichment for ion channel genes, ionotropic receptors, GPCRs, post-synaptic scaffolding and signaling components, and homeodomain TFs, but not other TF families, such as C2H2 zinc finger TFs (**Figure 4B, Supplemental Table 17**). In comparison to the conserved pan-neuronal genes, genes in these families, as well as neurotransmitter pathway genes, tend to show highly specific expression to a small portion of neuronal subtypes in a given species, but altogether, essentially every neuron in every species expresses at least one member of each family (i.e. at least one class 1 ion channel, at least one cationic glutamatergic receptor, etc.) (**Figure 4C**).

**Figure 4:**
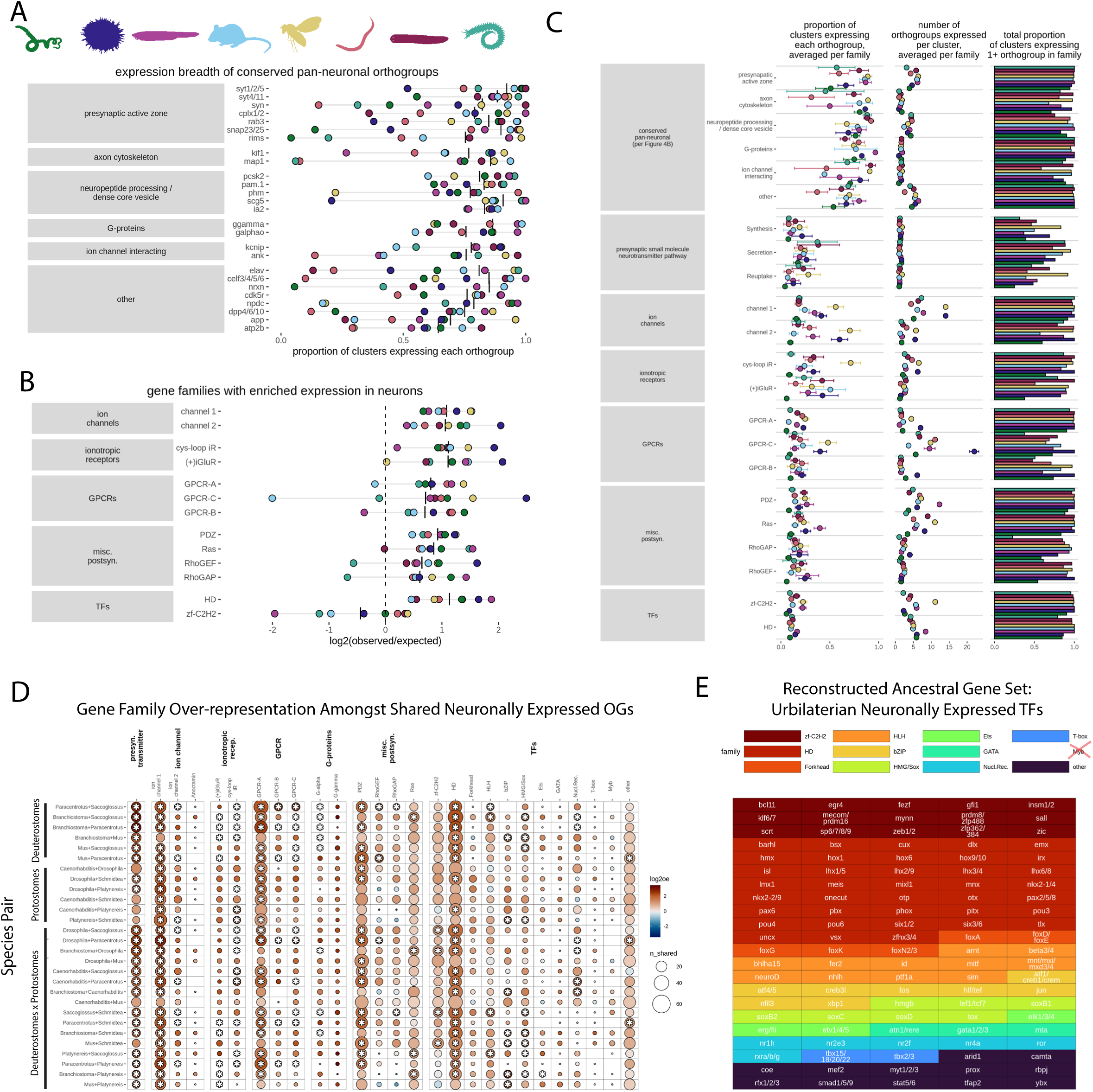
Cross-species comparisons reveal a conserved toolkit of pan-neuronal and neuron subtype identity genes. A) Top genes with high average expression breadth across species. Genes are categorized by published function. B) Select Pfam-defined gene families with high average enrichment of neuronal expression across species. zf-C2H2 TFs, which lack conserved enrichment, are also shown for comparison. C) Cross-species comparisons of expression breadth of conserved neuronally-enriched families. Left: Boxplot showing average proportion of cell clusters (x-axis) expressing genes in each orthogroup, orthogroups grouped by family and function (y-axis). Right: Bar plot showing total proportion of cell clusters (x-axis) expressing at least one orthologous gene in each family. D) Gene families enriched amongst genes with conserved neuronal expression. Pairwise comparisons amongst species (y-axis) for gene families (x-axis), including manually annotated presynaptic neurotransmitter pathway genes or Pfam-defined gene families. Asterisks indicate significant enrichment with adjusted p < 0.05. E) TFs expressed in neurons in the bilaterian LCA as reconstructed using a maximum parsimony approach (see Methods, **Supplemental Table 18**), with TFs colored by family. Note high representation of the HD family.

Further analysis reveals that the similarities amongst species extend beyond just the gene families expressed in neurons: in many cases, the exact same genes in these families are expressed in neurons. The gene families that are enriched in neurons within individual species are also enriched amongst the orthogroups that show conservation of neuronal expression across species (**Figure 4C-D**).The degree of similarity between each species’ “neuronal gene toolkits” (including TF set and specific effector sets) is largely determined by evolutionary relatedness and not by overall nervous system architecture (centralized or net) or lifestyle (terrestrial or aquatic) (**Supplemental Figure 6**), whereas differences are due to the origin of new genes as well as recruitment or loss of neuronal expression of ancient genes (**Supplemental Figure 6, Supplemental Table 18**). Based on cross-species similarities, we can reconstruct sets of ion channels, ionotropic receptors, GPCRs, and TFs expressed in neurons in the bilaterian last common ancestor (**Figure 4E**, **Supplemental Table 18**). The ancestral TF set shows a high representation of homeodomains: of the 110 predicted urbilaterian neuronally-expressed TFs, 38 (over one-third) are homeodomains (**Figure 4E**). This includes all 6 of the urbilaterian LIM-HD genes (*isl*, *lhx1/5*, *lhx2/9*, *lhx3/4*, *lhx6/8*, and *lmx1*) and 3 out of 5 urbilaterian POU-HD genes (*pou3*, *pou4*, and *pou6*, but not *pou1* or *pou2*) (J. F. Ryan et al. 2006). The highly complementary and altogether broad pattern of expression of LIM-HDs is conserved across most of the species (**Supplemental Figure 7**). Thus, despite the stark differences in form and function of the nervous systems of our comparison lineages, we found significant overlap in neuronal gene expression, with these genes following two entirely distinct patterns: 1) broadly pan-neural genes which impart a generic neuron identity and 2) neural identity genes which are found enriched in just a small fraction of the neurons in a species and impart a specific functional identity, including an ancient TF repertoire enriched for HDs

We next set out to compare the patterns of usage of this core effector toolkit across species to assess whether there is evidence for conserved neuron subtypes. Recent comparisons of single-cell transcriptomic data between distantly related species have relied on SAMap, which excels in identifying similarities between broad cell classes ((Tarashansky et al. 2021; Robertson et al. 2024; Piovani et al. 2023) and **Supplemental Figure 3B**). However, in the absence of strong one-to-one or few-to-few relationships between cell populations from different species, SAMap integrations can be difficult to interpret: strong interspecies hits can be driven by a small number of genes (Scully et al. 2025). Thus, we utilized SAMap as one tool to identify cross-species transcriptomic similarities, and sought an alternate method as a primary means of comparison. Based on the success of gene set and gene expression specificity-based approaches (Tosches et al. 2018; Shafer et al. 2022; Scully et al. 2025), we compared groups of cells (whole body clusters or neuronal subclusters) within and across species by comparing the sets of genes expressed in each subcluster, while more heavily weighing genes with a more restricted expression breadth (see Methods). When comparing whole body clusters between zebrafish, fly, and the hemichorate, this approach (**Supplemental Figure 8A**) recovered many of the same class-level similarities as SAMap (**Supplemental Figure 3B)**, while also identifying related cell populations within species (**Supplemental Figure 8B**), confirming its utility.

When we used the specificity-weighted set approach to compare neuronal subclusters, we detected similarities within known neuron “families” within species, for example, hypothalamo-prethalamic primordium (HyPTh) neurons in *Branchiostoma* and dorsal root ganglion neurons, olfactory bulb neurons, and spinal cord interneurons in mouse (**Figure 5A**, red labels 1, 2, 3, and 4). However, this method did not return strong similarities between particular neuron subpopulations between phyla (**Figure 5A**). SAMap identified a small number of strong subcluster alignments (**Supplemental Figure 9A**), but the sets of genes driving these similarities (**Supplemental Table 19**) tend to be small, or include many broadly expressed genes, or be enriched for ribosomal proteins (**Supplemental Figure 9B**), indicating that these hits are driven by only weak similarities, metabolic state, or cell quality. There is one set of exceptions, a set of subclusters with high similarity that can be identified by both the specificity-weighted set method and by SAMap. These subclusters includes the *Platynereis*, *Schmidtea,* and *Branchiostoma* populations sharing expression of ciliation genes (**Figure 5A**, red label 5, and **Supplemental Figure 9B**); the signature uniting these populations were previously observed via within-species analysis of the *Schmidtea* data (Fincher et al. 2018).

**Figure 5:**
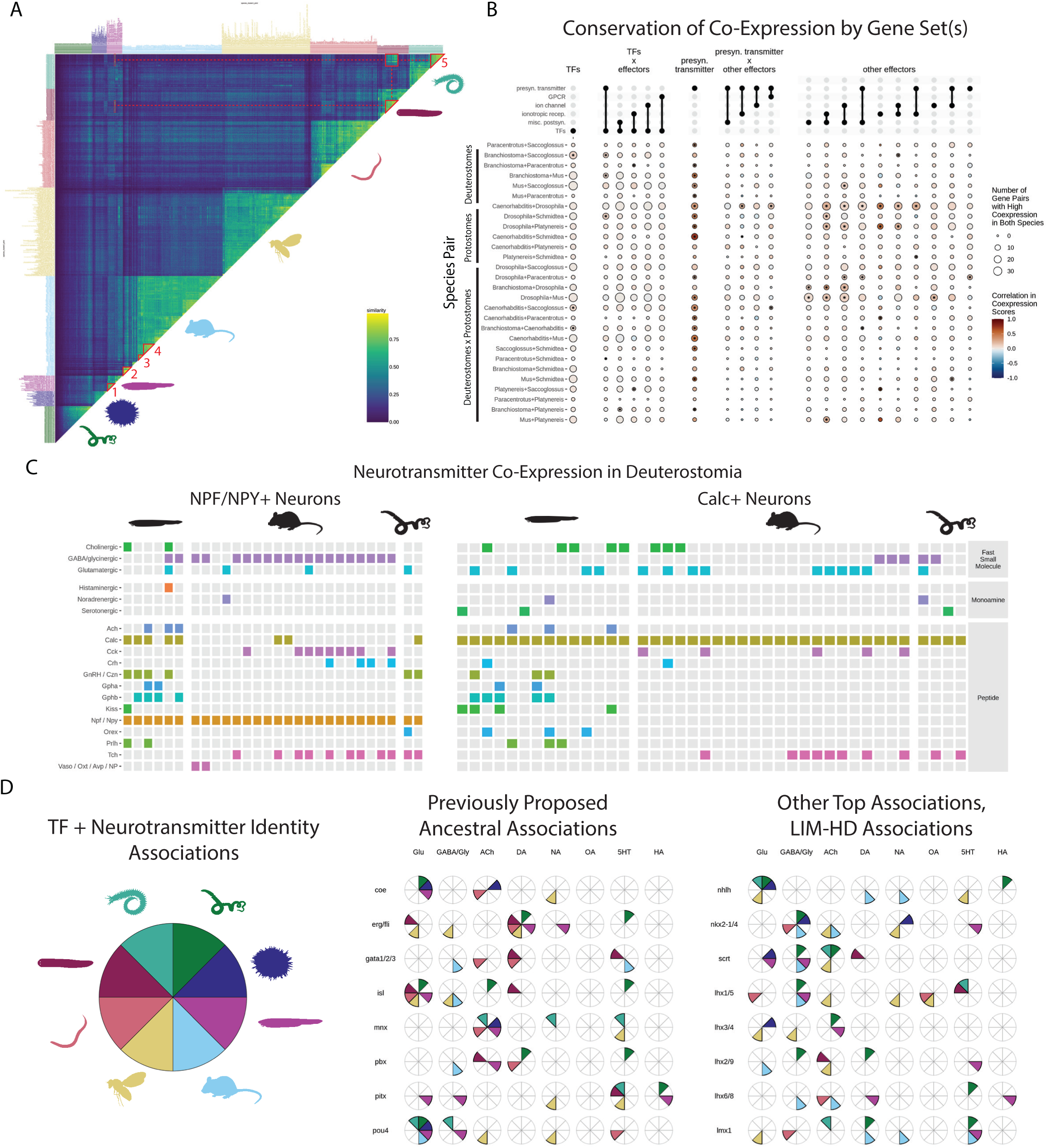
Widespread evolutionary turnover in the patterns of co-expression of ancient neuronal genes. A). Global transcriptional similarity between neuronal subtypes within and between species. B). Cross-species comparison of patterns of co-expression amongst gene sets. Co-expression correlation and number of highly co-expressed genes. C). Evolutionary turnover in the patterns of co-transmission of small molecules and peptides. D) Conservation of significant TF-neurotransmitter identity associations. Filled slices in pie charts indicate detection of association in the corresponding species, as shown in the key (left). Center shows associations for TFs previously proposed to have conserved roles in specifying particular neurotransmitter identities: *coe* and cholinergic identity (Popsuj and Stolfi 2021; Catela et al. 2019), *erg/fli* and serotonergic identity (Lloret-Fernández et al. 2018) or dopaminergic identity (Flames and Hobert 2009), *gata1/2/3* and serotonergic identity (Lloret-Fernández et al. 2018), *isl* and glutamatergic identity (Nomaksteinsky et al. 2013), *mnx* and cholinergic identity (Nomaksteinsky et al. 2013), *pbx* and dopaminergic identity (Remesal et al. 2020), *pitx* and GABAergic identity (Westmoreland et al. 2001), and *pou4* and glutamatergic identity (Nomaksteinsky et al. 2013). Right shows most conserved TF-neurotransmitter associations, as well as LIM-HD-neurotransmitter associations. Note limited conservation of most associations.

To rule out that examining global similarity did not result in our overlooking more specific conservation of the genes most critical for neural identity and function, we attempted restricting our analysis to just genes known to contribute to key aspects of the neuronal phenotype–neurotransmitters, ion channels, transmembrane receptors, over-represented families (**Supplemental Figure 10**). When comparing subclusters using only small molecule neurotransmitter-related genes, we identify strong cross-species similarities amongst subsets of subclusters (**Supplemental Figure 10A**), driven by the conservation of the patterns of gene co-and contra- expression underlying the small molecule neurotransmitter pathways (**Supplemental Figure 10B**). In contrast, other sets of effector or TFs do not recover high cross-species similarities (**Supplemental Figure 10C-D**). Overall, these results indicate that there are few, if any, cases of neuronal transcriptomic identity being conserved between distantly related bilaterian taxa.

As comparisons of global transcriptional similarity can mask signals of conserved gene co-usage, we next set out to explicitly quantify the conservation of gene co-expression across species. Specifically, we used a jaccard-based approach to quantify co-expression amongst orthogroups within species, then compared the overall similarity in co-expression patterns for sets of orthologous genes across species. As expected, neurotransmitter pathway genes show strong conservation of patterns of co-expression (**Figure 5B**, “pre-syn. transmitter”). In contrast, the patterns of co-expression within ion channels, GPCR, and fast post-synapse gene sets show limited enriched similarity across species (**Figure 5B**). Moreover, these sets of genes are not co-expressed with each other in strongly conserved patterns (**Figure 5B**). While there are dozens of instances of orthogroup pairs showing high co-expression across two or more species (**Supplemental Table 20**), overall, the patterns of effector gene co-expression is not highly conserved across different bilaterian phyla.

It has been suggested that conservation of regulatory gene cooperativity may be under constraint, even in the absence of coupling to conserved effector genes (Wagner 2014). Moreover, conserved associations between sets of TFs and certain neuron cell phenotypes have been proposed to be conserved since the bilaterian last common ancestor (Nomaksteinsky et al. 2013; Lloret-Fernández et al. 2018; Remesal et al. 2020; Flames and Hobert 2009; Westmoreland et al. 2001; Catela et al. 2019; Popsuj and Stolfi 2021; Erclik et al. 2009; Arendt 2003; Valencia et al. 2021; Arendt et al. 2004; Paganos et al. 2022; Denes et al. 2007; Tessmar-Raible et al. 2007). Thus, we next examined whether there was any evidence for conserved TF programs among phyla or conserved association between specific TFs and neuronal effector genes. As in our comparisons amongst effector genes, we find limited conservation of patterns of TF-TF co-expression, and limited conservation of the patterns of co-expression between TFs and effectors (**Figure 5B**). Overall, these results suggest that there are few deeply and broadly conserved neuronal gene expression modules beyond the minimal set of genes required to produce, process and transport the neurotransmitter used by the cell. Rather, the same TF and effector genes are co-expressed in distinct patterns in distantly related species–indicative of “remixing” of effector genes following the divergence of these species.

The outcomes of this evolutionary reshuffling can be observed by comparing, for example, the patterns of small molecule plus neuropeptide co-transmission across deuterostome species. We found that homologous neuropeptides (see Methods and **Supplemental Table 7**) show very different patterns of co-expression in *Branchiostoma*, *Mus*, and *Saccoglossus* (**Figure 5C**), indicating that, while small molecule plus peptide co-transmission is common and even involves many of the same transmitters across species, the combinations of molecules that are released from the same neuron are frequently different in different species; co-transmission identity is not evolutionarily stable.

Instances of ancient TF-transmitter phenotype associations have been previously proposed (Nomaksteinsky et al. 2013; Lloret-Fernández et al. 2018; Remesal et al. 2020; Flames and Hobert 2009; Westmoreland et al. 2001; Catela et al. 2019; Popsuj and Stolfi 2021). Our analysis indicates that the patterns of co-expression between TFs and transmitter pathway effector genes are generally not any more similar across species than would be expected by chance (**Figure 5B**). Previously proposed conserved TF-transmitter associations (**Figure 5D**, center, associations with black borders) can be identified in a few species, but most are recovered in fewer than half of species (**Figure 5D**). For example, *pbx* is significantly associated with dopaminergic identity (Remesal et al. 2020) in only two species; the same is true for the *coe*-cholinergic association (Popsuj and Stolfi 2021; Catela et al. 2019). Amongst the few TFs with strong signal of conserved roles in neurotransmitter identity are some previously proposed candidates, such as *erg*/*fli* with dopaminergic identity (Flames and Hobert 2009) and *isl* and *pou4* with glutamatergic identity (Nomaksteinsky et al. 2013) (**Figure 5D**, center), as well as associations between *nkx2-1/4* and GABA/glycinergic identity and *nhlh* and glutamatergic identity (**Figure 5D**, right). Still, even these top associations are only found in a subset of species, indicating that these potentially ancestral associations were lost in some lineages. The LIM-neurotransmitter associations we observed in *Saccoglossus* are not broadly conserved (**Figure 5D**, right); in general, TF-neurotransmitter identity associations are highly species-specific (**Figure 5D, Supplemental Figure 11**). Overall, the limited cases of conservation suggest an evolutionary history involving many instances of losses and independent (sometimes convergent) gains of TF -neurotransmitter associations. Thus, in strong contrast to previous anecdotal observations and general assumptions, the associations between TFs and neurotransmitter identity are largely lineage-specific.

## Discussion

Here, we generate a whole-body single cell transcriptome atlas for a hemichordate worm, a phylogenetically and historically important lineage for our understanding of nervous system evolution. Within the relative simplicity of the acorn worm nerve net, with its loosely condensed cords and vast peripheral plexus, we detect a great diversity of over 70 neural cell types. We find, despite its largely non-centralized structure, the hemichordate nervous system utilizes the same peptidergic, excitatory, inhibitory and monoaminergic transmitters and similar sets of ion channels and ionotropic receptors as centralized bilaterian nervous systems. Of the many distinct neurons we characterize, it is notable that all express the canonical neuropeptide processing genes needed to cleave, amidate and package neuropeptide transmitters for release. From an evolutionary perspective, the pervasive nature of peptidergic signaling also aligns with its likely role as the dominant neurotransmitter in early nervous systems. Previous work in cnidarian nerve nets (Plessier and Marlow 2026; Gründer and Assmann 2015; Morris Little et al. 2025) has shown that these systems are largely peptidergic in nature. Combined with molecular studies that show synaptic-like vesicles and neuropeptide-based signaling in choanoflagellates and placozoans (Yañez-Guerra et al. 2022; Göhde et al. 2021; Varoqueaux et al. 2018; Smith et al. 2014; W. Li et al. 2025; Najle et al. 2023), it is likely that neuropeptide transmission is the ancestral state for animal neurons.

In addition, nearly all hemichordate neurons show signatures for co-transmission of multiple neuropeptides or neuropeptides alongside small molecules. Hemichordate neurons are not alone in this propensity: recent systems-level characterizations of vertebrate and nematode neurons have revealed that the majority of neurons display co-transmission of small molecules and peptides (Z. Yao et al. 2023; Taylor et al. 2021). It has been suggested that co-transmission may provide opportunities for neurons to modulate their communication with synaptic partners or transmit multi-layered signals. Thus, despite the morphological simplicity and modest number of distinct neuron types in the hemichordate nervous system, the potential functional diversity of neuronal signaling conveyed through such a system could be quite high.

Bilaterian neurons have long been considered a homologous cell type based on morphological and functional similarities, as well as molecular features such as occurrence of notable pan-neural markers such as ELAV, synapsin and synaptotagmins. By examining the basis of neural identity in hemichordate and five other distant bilaterian phyla, we uncover a robust cross-species pan-neural signal based primarily on shared usage of the pre-synaptic active zone genes and widespread expression of the neuropeptide processing machinery, suggesting that these features unite bilaterian neurons.

In contrast to the robust pan-neuronal signature we identify, we detect, using several distinct methods including a highly permissive pair co-expression analysis, no signal for cross-phyletic neural subtype conservation. However, we observe expression of at least 8 small molecule neurotransmitter pathways in separate neuron populations across protostomes and deuterostomes alike. This data, in combination with the large size of the reconstructed ancestral gene toolkits of ion channels, receptors, and TFs, raise the possibility that the Urbilaterian possessed as many as dozens of distinct neuron subtypes. But we find no evidence that any ancestral subtype-specific gene expression programs that may have existed were conserved over geologic timescales (Figure 6, top); rather, the history of the bilaterian nervous system was defined by a pattern of adapting preexisting neuron diversity through changes in gene co-expression patterns (Figure 6, bottom). Indeed, recent comparative work has illustrated changes in the expression of orthologous genes in evolutionarily equivalent neuron populations amongst relatively closely related (co-phyletic) species (Hain et al. 2022; Shafer et al. 2022). Even in the genus *Caenorhabditis*, which has diversified so recently that each neuronal subtype in one species shows a single obvious molecular equivalent in each other species, there are still dozens of genes, including ion channels, neurotransmitter receptors, and neuropeptides, that show qualitative cross-species differences in which neurons they are expressed within (Toker et al. 2025). An extrapolation of these patterns over hundreds of millions of years would seem consistent with the major differences in co-expression of ancient neuronal genes that we observe when making comparisons across phyla.

**Figure 6:**
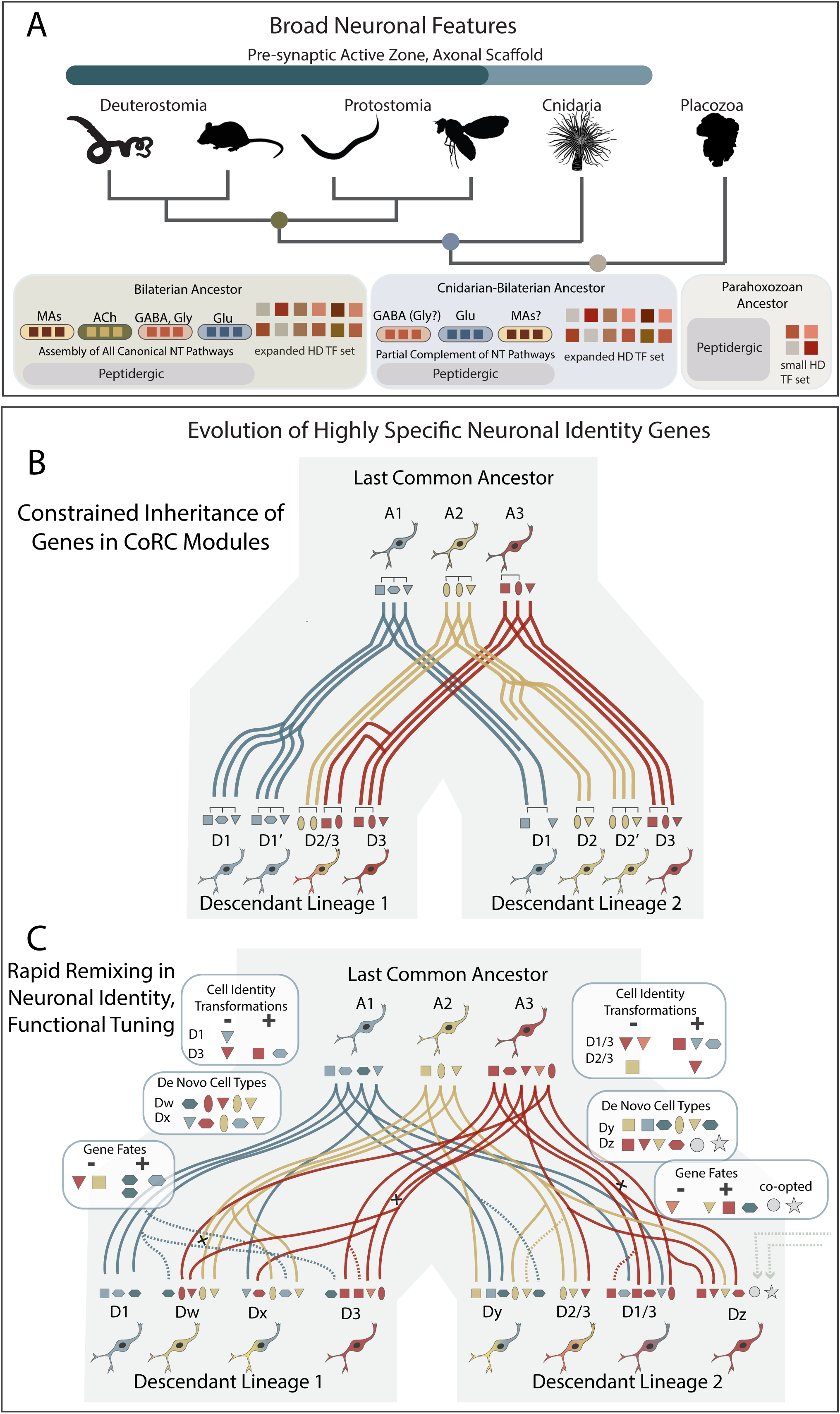
Evolutionary scenario for the evolution of neuronal features. A) Evolution of broadly neuronal molecular features in neurons. Homeodomain TFs (HD TFs) and discrete peptidergic cell types are inferred to be present in the LCA of Placozoa and Cnidaria+Bilateria. Diversification of HD TFs has occurred prior to the divergence of the LCA of Cnidaria and Bilateria. GABA and glutamate pathways are utilized as transmitters in neurons. Some evidence supports a possible partial evolution of monoamine (MA) pathways. The last common bilaterian ancestor (LCBA) has evolved a full complement of fast transmitter, MA and peptidergic neurotransmitter pathways. B) Model of cell type evolution considering limited re-mixing of genes which are constrained by regulation in co-regulatory complexes. Genes which are regulated by largely overlapping TFs are maintained as co-expressed modules, thus limiting the amount of de novo transcriptome assembly and novel cell type formation. C) A model in which very limited co-regulation of genes largely allows for independent sorting of components of the transcriptome of an ancestral cell type (A) to new cell types in descendant lineages (D). Thus, composite, *de novo* and highly divergent resultant cell types are to be expected.

Still, changes in gene co-expression alone are insufficient to explain the early evolution of bilaterian neurons, in particular when considering the massive molecular diversity of neurons present in lineages with highly centralized nervous systems. Evidently, changes in gene co-expression occurred alongside lineage-specific expansions to the number of unique neuronal subtypes. As new subtypes emerged independently, the types of genes used to build these distinct identities were the same between phyla and include largely the same families of GPCRs, ion channels, and post-synaptic scaffold proteins. This rapid reshuffling of neuronal effector genes has occurred alongside previously recognized rampant gene duplication in these families (Hehmeyer et al. 2024; Jin et al. 2023; Fernández and Gabaldón 2020), suggesting a larger pattern in which the adaptation of animal nervous systems to novel environments, behaviors, or body plans occurs through a high turnover and diversification of neuron subtypes.

Given the lack of specific similarity amongst neuron subtypes from different phyla, it is conceivable to envision a wide variety of regulatory programs could have been deployed to specify a diversity of terminally differentiated neurons. Yet, we find a strong signal for an ancestral bilaterian neuronal TF lexicon highly enriched for HD transcription factors. In each lineage examined, there is again, a strong signal for reshuffling with little evidence for the coupling of specific TFs with distinct classes of neurons across phyla. We propose that a code of heterodimerizing TFs, such as the HDs, provide an ideal substrate to generate combinatorial complexity of the transcriptional code and are ideally suited to specifying cellular identities amongst the wide diversity of distinct neuron types found within each lineage. While most of the homeodomains we observe show highly restricted and specific expression, a notable exception are the LIM-HD genes which are expressed across broad groups of neuronal subclusters. It has been previously suggested in flies, mice, and sea anemones that LIM-HDs may establish a code specifying aspects of neural identity such as connectivity or neurotransmitter usage (Srivastava et al. 2010; Thor et al. 1999; Allen et al. 2020; Z. Yao et al. 2023). It remains to be established whether the LIM-HDs contribute to features of a pan-neuronal or subtype-specific code more broadly in animals. Strikingly, previous phylogenetic analyses have suggested that the diversification of HD genes, including the LIM-HD genes, occurred in the lineage preceding the last common ancestor of Cnidaria and Bilateria (Joseph F. Ryan et al. 2010; J. F. Ryan et al. 2006). Thus, it would appear that their diversification closely predates, if not parallels, the diversification of neuronal subtypes (Hobert 2021).

While the building blocks which are used to generate these unique neural identity codes in Bilateria are conserved, we find that their co-expression largely is not – our findings stand in stark contrast to existing models of how cell-type specific regulatory logic may evolve (Wagner 2014; Arendt et al. 2016, 2019) (Figure 6B). These models propose or presuppose deep conservation of the regulatory and effector programs associated with terminal differentiation and the maintenance of mature cell identity, and, indeed, some comparative studies do indicate that certain neural TF+effector modules can be maintained over great evolutionary timescales, at least within phyla (Jin et al. 2023; Tommasini et al. 2025; Hahn et al. 2023; Junqiang Wang et al. 2024). However, our results more strongly evoke earlier models of gene regulatory network evolution, that proposed high flexibility of the most "peripheral" components of the developmental network–the circuits responsible for generating the effector programs of the terminally differentiated cell types–in contrast to the greater conservation of earlier-acting network components (Davidson and Erwin 2006) (Figure 6C). Indeed, our observations of turnover of gene expression contrasts the strong pattern of conservation (albeit, based on candidate gene approaches) seen in the early patterning of the deuterostome neuroectoderm (Christopher J. Lowe et al. 2003; Y. Yao et al. 2016; Pani et al. 2012; Lin et al. 2025; Benito-Gutiérrez et al. 2021) and in the neuroprogenitor and immature neuron programs shared across even greater evolutionary distances (Allan and Thor 2015; Slota and McClay 2018; Richards and Rentzsch 2015). Increased phylogenetic and ontogenetic sampling both within and between phyla will be necessary to definitively resolve neural origins and fully elucidate the patterns of evolution of the neural gene regulatory network. Still, the findings here necessitate a re-evaluation of our current assumptions for widespread conservation of neural subtypes in Bilateria.

## Acknowledgements

We would like to thank Sebastien Bastide for assistance with scRNA-seq library training and for constructive feedback on project design and analysis. We thank members of the Marlow lab for many useful discussions and insightful feedback. We would also like to thank members of the Lowe and Range Labs for assistance with field collections and culture maintenance. We thank Urs Schmidt-Ott and Victoria Prince for their thoughtful feedback on initial drafts of the manuscript. We thank the Marine Biological Laboratory for supporting this work via the Whitman Fellows Program and access to imaging facilities and other critical infrastructure. We also acknowledge the Organismal Biology and Anatomy Light Microscopy Imaging Facility. We thank the National Institutes of Health for funding through award 5R35GM155159 to HM. We also acknowledge NIH T32 training grant GM139782 for support of JH.

## Methods

### Hemichordate animal culture and scRNA-seq

Adult *Saccoglossus kowalevskii* worms were collected from Waquoit Bay, Massachusetts in September 2022 and 2023. Spawning of female adult worms, fertilization of eggs, and culture of embryos were carried out as previously described (Christopher J. Lowe et al. 2004). Hatched animals were either sampled unfed (1-gill-stage) or fed Reef Nutrition® Phyto-Feast® Live (Reed Mariculture Inc., Campbell, CA, USA) daily for several weeks until they reached the 3-gill stage, then starved for 4 days prior. For single cell sequencing experiments, young 1-gill-stage or 3-gill-stage animals were dissociated into a single cell suspension by gently titurating in a dissociation medium comprised of 0.9X calcium and magnesium -free seawater (0.9X CMFSW: 445mM NaCl, 8.7mM KCl, 24.8mM NaHCO3, 45mM Tris-HCl pH8) and 0.25mg/mL LiberaseTM for 20 - 90 minutes, depending on stage. Following dissociation, cell viability was confirmed by staining with Calcein AM (Invitrogen C1430), Hoechst, and Propidium Iodide (Invitrogen P3566), and the cell concentration was adjusted to target a total yield of 20K cells. For the 1gill stage sample, 43.2uL of cell suspension was directly mixed with master mix (18.8 uL of RT Reagent B (10X Genomics #2000165), 8.7uL of RT Enzyme C (10X Genomics #2000102), 2.4 uL of Template Switch Oligo (10X Genomics #3000228), and 2.0 uL of Reducing Agent B (10X Genomics #2000087)) just prior to droplet encapsulation using the 10X Chromium Controller. For the 3gill stage sample, we modified this step to incorporate a strategy similar to one recently reported (Scully and Klein 2025) to reduce the hypo-osmotic lysis of cells: 31.8 uL of master mix was diluted with 21.6 uL of mannitol-supplemented 1X CMFSW (0.7M D-Mannitol, 495mM NaCl, 9.7mM KCl, 27.6mM NaHCO3, 50mM Tris-HCl pH8) prior to the addition of 21.6 uL of 2X-concentrated cell suspension. Libraries were generated from single cell emulsions using the 10X Genomics scRNA-seq v3.1 kit and sequenced on the NextSeq 550 platform.

### Generation of improved gene models for *Saccoglossus kowalevskii*

*Saccoglossus kowalevskii* NCBI gene models (RefSeq Annotation GCF_000003605.2) and JGI gene models ((Simakov et al. 2015), accessed at https://groups.oist.jp/molgenu) were first lifted over to the chromosome-scale genome (Bastide et al., in press) using liftoff (Shumate and Salzberg 2021). Then, gffcompare (Pertea and Pertea 2020) was used to compare transcripts; redundant transcripts were removed and genes with overlapping exons were combined. This approach resulted in a single-copy proteome with a BUSCO score of 96.7% (S:93.9%,D:2.8%,F:2.3%,M:1.0%) when run using database “metazoa_odb10” (n=954). cDNA reads from all samples were mapped using hisat2 (Kim et al. 2019) for a 3’ extension of up to twice the median gene length using geneext (Zolotarov et al. 2026). Mitochondrial genome and gene models from NCBI (NC_007438.1) were added to generate a combined nuclear and organelle gene model set.

### Generation of improved gene models for *Branchiostoma floridae*

*Branchiostoma floridae* NCBI gene models (RefSeq GCF_000003815.2) were lifted over to the Li Lab genome ((Dai et al. 2024), accessed at doi.org/10.57760/sciencedb.08801) using liftoff. Then, gffcompare was used to compare NCBI transcripts to those from the Li Lab gene models; perfectly redundant transcripts were removed and transcripts with overlapping exons were combined into the same gene. This approach resulted in a single-copy proteome with a BUSCO score of 98.0% (S:96.0%, D:2.0%, F:0.2%, M:1.8%), greater than the scores for the NCBI and Li Lab proteomes. cDNA reads from T1-stage scRNA-seq samples (Dai et al. 2024) were mapped using hisat2 for a 3’ extension of up to the median gene length using geneext. Mitochondrial genome and gene models from NCBI (NC_000834.1) were added to generate a combined nuclear and organelle gene model set.

### Generation of improved gene models for *Platynereis dumerilii*

Because we observed that the BUSCO score of *Platynereis* gene models dropped significantly when generating a single-copy proteome, a new set of gene models was generated with reduced chimeric genes. Specifically, we generated gene models for a scaffold-level genome (NCBI accession GCA_026936325.1) from publicly available short and long read RNA-seq data (Conzelmann et al. 2013; H.-C. Chou et al. 2016; Vellutini et al. 2024; Schenk et al. 2019; Paré et al. 2023) using hisat2 and stringtie (Kovaka et al. 2019). We then applied mikado (Venturini et al. 2018) to these custom gene models plus gene models hosted by ENSEMBL (https://metazoa.ensembl.org/Platynereis_dumerilii_GCA_026936325.1cm/Info/Annotation) to split chimeric genes and select amongst transcripts. This approach resulted in a single-copy proteome with a BUSCO score of 92.9% (S:90.1%, D:2.8%, F:0.7%, M:6.4%), greater than the ENSEMBL score. Reads from all scRNA-seq samples (Milivojev et al. 2025; Stockinger et al. 2024) were mapped using hisat2 for a 3’ extension of up to the median gene length using geneext. Mitochondrial genome and gene models from NCBI (NC_000931.1) were added to generate a combined nuclear and organelle gene model set.

### scRNA-seq data mapping and processing

We generated new multi-sample scRNA-seq datasets from previous and newly-generated data from *Saccoglossus kowalevski* and from publicly available data from *Platynereis dumerilii* (Stockinger et al. 2024; Milivojev et al. 2025) and *Branchiostoma floridae* (Dai et al. 2024). Single cell reads were demutiplexed, mapped, and quantified using STARSolo (Kaminow et al. 2021). Total, unspliced, and spliced read counts were quantified by applying the default (“gene”), and “velocyto” quantification methods. Soup and doublets were removed using decontX and doubletFinder, respectively. We found that a variety of scRNA-seq datasets from whole-body or multi-tissue invertebrate samples include a large number of droplets with a disproportionately low fraction of unspliced reads relative to their total read count; these droplets contain cytoplasmic (anucleated) debris rather than quality, intact cells (DropletQC paper, other paper). After implementing a step to remove cells with an unspliced-to-total counts ratio below a per-dataset or per-cluster cutoff, we integrated samples using Harmony while regressing out nuclear fraction and mitochondrial content. Harmony was reapplied to subsets of cells to generate sub-clusterings (i.e. neurons). For *Saccoglossus* and *Platynereis*, we used uncorrected counts as the default assay to avoid overcorrection. We annotated *Saccoglossus* clusters as described in the text and according to the list of known or functionally-informative marker genes, as provided in **Supplemental Table 1**.

We recreated the *Platynereis* and *Branchiostoma* annotations from original source publications by using available annotations from the same cell sets, when available, otherwise by comparing expression of expert annotated marker genes. For *Branchiostoma*, we labeled glial cells (located within the previously-annotated neural clusters) based on published marker genes (Bozzo et al. 2021) and removed them from the neuronal analysis. Additionally, we removed cycling neural populations (progenitors) based on expression of cell cycle signature genes (see below).

For *Platynereis*, our neuronal subclustering represents the first dataset of its kind for an annelid. We validated our neuronal subclustering by assessing the expression of small molecule neurotransmitter and neuropeptide genes, which have previously been demonstrated to have distinct, restricted expression patterns in the nervous system of this animal.

For other species, publicly available data (Paganos et al. 2025; Zeisel et al. 2018; D. Lee et al. 2025; H. Li et al. 2022; Davie et al. 2018; Allen et al. 2020; Özel et al. 2021; Hopkins et al. 2023; Taylor et al. 2021; Fincher et al. 2018) were used without remapping or full preprocessing. Partial preprocessing was carried out for *Mus* and *Drosophila* data as follows. For *Mus*, we identified and removed neuron progenitor populations based on original dataset annotations. For *Drosophila*, combining several neuronal datasets required standardizing genes and then integrating data. First, we lifted over all gene symbols to match those present in the annotation from Flybase Release FB2024_06 based on stable “FBgn” IDs. Then, neurons and non-neurons were distinguished within each dataset based on the provided source annotations or based on the expression of non-neuronal cell markers as identified in the original publications. All neurons were integrated into a unified set of clusters using Harmony. We confirmed that equivalent populations from different datasets fell within the same clusters.

### Gene orthology, functional annotation, and ontology

Single-copy proteomes were generated by selecting the longest peptide sequence per gene. For manually aggregated gene model sets (hemichordate, amphioxus, and zebrafish), peptides from all original sources were considered regardless of the source of the longest transcript. Then, we grouped genes in these protomes into orthogroups using an extended orthology analysis built around OrthoFinder. First, we ran OrthoFinder version 3.1 (Emms et al. 2025) using single copy proteomes from 13 species that passed our strict quality and relatedness requirements, including 6 out of 9 species used for scRNA-seq data comparisons (*Saccoglossus*, *Branchiostoma*, *Mus*, *Danio*, *Drosophila*, *Platynereis*), as well as 7 additional species included to serve as phylogenetic intermediate and additional sampling. We did not include proteomes from species in our scRNA-seq comparisons that negatively impacted OrthoFinder results. These are the proteome from *Paracentrotus*, which shows a lower BUSCO score than other species (87.9%, S:86.6%, D:1.4%, F:8.8%, M:3.2%); the proteome of *Caenorhabditis*, a member of Nematoda, which has experienced a higher rate of coding sequence evolution compared to other bilaterian phyla (Laumer et al. 2019); and the transcriptome-based, and thus redundant, proteome from *Schmidtea*, another member of another phylum with a high rate of coding sequence evolution, Platyhelminthes (Laumer et al. 2019). To add these species to our orthology analysis, we ran blastp using the single-copy (*Paracentrotus*, *Caenorhabditis*) or redundant (*Schmidtea*) proteomes from these species as queries against the combined proteomes from all other species, and added each peptide to the orthogroup of the gene with the top hit with an e-value of less than 1e-4. The final orthology results are provided in **Supplemental Table 15**.

Cell cycle genes were taken from https://github.com/hbc/tinyatlas and carried over to other species based on the orthology analysis results. Cell cycle genes were validated by assessing co-expression, and genes not showing strong signatures of co-expression with other cell cycle genes were removed.

GO terms and Pfam domains were annotated for every proteome using Eggnog Mapper. TF genes and their families identified using the jaspar inference tool (Fornes et al. 2020), except that three Pfam domains generally not associated with TF activity (RMP, mTERF, and CENP-B) were excluded from the TF prediction. Homeodomain classes were annotated by using BLAST to compare all homeodomain-containing genes to the the HbxFinder vertebrate homeodomain databases (Mulhair et al. 2023; Zhong and Holland 2011) (https://github.com/PeterMulhair/HbxFinder/). All non-TF genes with TF-type DBDs (some chromatin modifiers and general transcriptional machinery components) were excluded from the ancestral state reconstruction in Figure 4E.

Neuropeptide identification and homology relations were based on previously published results (Tian et al. 2016; Semmens et al. 2016; Andrade López et al. 2023; Mirabeau and Joly 2013; Jékely 2013; Yañez-Guerra et al. 2022, 2020; Yañez Guerra and Zandawala 2023; Elphick 2010; Tinoco et al. 2021; Veenstra 2021; Schiffer et al. 2024; Zieger et al. 2021; Zandawala et al. 2017; Theofanopoulou et al. 2021; Tando and Kubokawa 2009a, 2009b; Huang et al. 2025; On et al. 2022) (**Supplemental Table 7**) with the exception of one new *Saccoglossus* glycoprotein hormone beta subunit gene, which we discovered through our orthology analysis (**Supplemental Table 15**).

### Single-species and comparative analysis of scRNA-seq data and associated gene sets

Throughout our analyses, p-values were corrected using the False Discovery Rate (FDR) method, and results with corrected p-values of less than 0.05 were considered significant. Prior to differential gene expression analysis we corrected for differences in sequencing depth using either Seurat::PrepSCTFindMarkers or batchelor::multiBatchNorm. We then identified positively differentially expressed genes by comparing expression between each cluster and all other cells (whole body data analysis) or between each neuron subcluster and all non-neuronal cells (neuron subcluster analysis) using the logistic regression (“LR”) method included in Seurat’s FindMarkers function, allowing us to remove batch effects by including batch as a latent variable. The genes identified as positively and significantly differentially expressed in a given neuronal subcluster were treated as the “marker” genes or “expressed” genes for that subcluster, as mentioned throughout the text.. Gene set enrichment and over-representation analysis was carried out using the fgsea (Korotkevich et al. 2021) R package.

For ancestral state characterization, we followed the dated species tree according to Carlisle et al. 2024. Ancestral state reconstructions were carried out using the phytools (Revell 2012) and phangorn (Schliep 2011) R packages. Reconstructions for quantitative traits (gene expression breadth) were carried out using a maximum likelihood -based approach, whereas reconstructions for binary traits (neuronal gene expression presence or absence) were carried out using maximum parsimony under an equal rates model. We took a conservative approach for the binary reconstructions, with all ambiguous states treated as lacking neuronal expression.

For SAMap interspecies clustering, we downsampled datasets to 40,000 cells maximum per species for whole body comparisons and 10,000 cells maximum per species for neuron vs. neuron comparisons, using a downsampling strategy that downsampled proportionally to size, such that no cells would be removed from the smallest cluster before cells were removed from any larger clusters. We then carried out SAMap while restricting gene homology to genes within the same orthogroup.

For cross-species comparisons of expression breadth, gene set similarity, and gene co-expression similarity, data from genes within the same orthogroup were combined within species.

Some previous work (Shafer et al. 2022; Scully et al. 2025) has shown that comparing binarized gene expression (above-basal expression or not) is an effective approach to control for interspecies differences in absolute expression levels, whilst other work (Tosches et al. 2018) has indicated that comparing the specificity of expression can improve accuracy: related cell populations (i.e. neuronal subtypes) may share expression of many of the same genes (ie pan-neuronal or broadly expressed neuronal genes) and evolutionarily-significant similarities and differences may only be detectable by focusing on genes with more restricted expression. Thus, for our within and cross-species cluster comparisons, we scored gene set similarity as follows. For a given cluster, *c*, in a given species, *s*, the expression of each orthogroup, *g*, binarized gene expression E*_g_*_,*s*,*c*_ is determined by our differential expression analysis, *K_s_* indicates the total number of clusters in species *s*, and *k_g_*_,*s*_ indicates the total number of clusters in species *s* for which E*_g_*_,*s*,*c*_=1. The expression specificity of each gene for each species and cluster is calculated as follows:

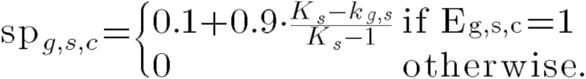

Thus, each gene expressed in a cluster has an expression specificity between 0.1 (gene is ubiquitously expressed across all clusters included in the comparison for a given species) and 1 (expression is entirely specific to the cluster in the given species). The minimum specificity value for an orthogroup, *m_g_*, is identified amongst the two clusters:

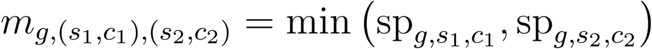

Only *m_g_* values greater than 0 are retained. The top *N* highest minimum specificity values across all orthogroups are used to calculate a similarity score:

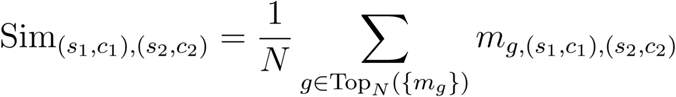

We used *N*=min(100, length(*m_g_*)) for whole transcriptome comparisons, and *N*=min(50, length(*m_g_*)) or *N*=min(10, length(*m_g_*)) for limited gene set comparisons (TFs only, ion channels only, neurotransmitters only, etc.). Scores fall between 0 and 1, with a score of 0 indicating the clusters do not share expression of any genes, while a score at or near 1 indicates that the clusters share all genes in common, with all genes showing highly specific expression.

Gene co-expression was measured using the jaccard test of (Chung et al. 2019), which calculates a normalized jaccard statistic that accounts for the expected set overlap relative to the number of data points (in this case, total number of clusters), and then calculates statistical significance using this statistic. Specifically, binarized expression values were compared across the total set of neuronal subclusters, allowing for expression co-occurence to be measured relative to expression breadth. Genes were considered “highly co-expressed" if they had positive and statistically significant normalized jaccard statistics. Co-expression of gene sets was compared across species using a spearman’s correlation test for the normalized jaccard statistics for pairs of genes.

Neurotransmitter identity was assigned based on expression of minimum identify-conferring gene sets as in **Supplemental Table 8**. Associations between TF genes and neurotransmitter identity were assessed using the same jaccard test as above.

### RNA fluorescent *in situ* hybridization chain reaction (HCR)

HCRs were carried out as described (Andrade López et al. 2023), with several modifications as also reported in Bastide et al. (in press). Specifically, samples were delipidated with 10% CHAPS in 1X PBS (Zhao et al. 2020) for 1 hour for 3-gill stage animals and 40 minutes for all 1-gill and embryo stage animals. Samples were then washed into 1X PBS, and photo-bleached with 15,000 lumen full-spectrum light at 4C for 2 hours. Following HCR and DAPI staining, embryos were stored in a dilute Optiprep-based solution (3.5M urea, 21% iodixanol, 0.1M NaCl, 0.5X PBS, 0.5X DAPI) at 4C for up to 24 hours. Embryos were then mounted in a high refractive index solution (7M urea, 42% iodixanol) (Hsu et al. 2022; Bedois et al. 2024) and imaged immediately. Tiled z-stack images were collected on a Zeiss LSM 900 confocal microscope at 20X resolution. When images were captured from only a small number of animals, additional, lower resolution, surface-level images were captured using a Zeiss Axio Imager M2 to serve as replicates. Raw image data was processed using Zen Blue and Fiji.

## Data Availability

New count matrices, gene models, processed R objects, and all code needed to reproduce these results are available on Zenodo (doi.org/10.5281/zenodo.19454501). Raw sequencing data will be made available on NCBI GEO at the time of publication.

---

**Supplemental Figure 1.**
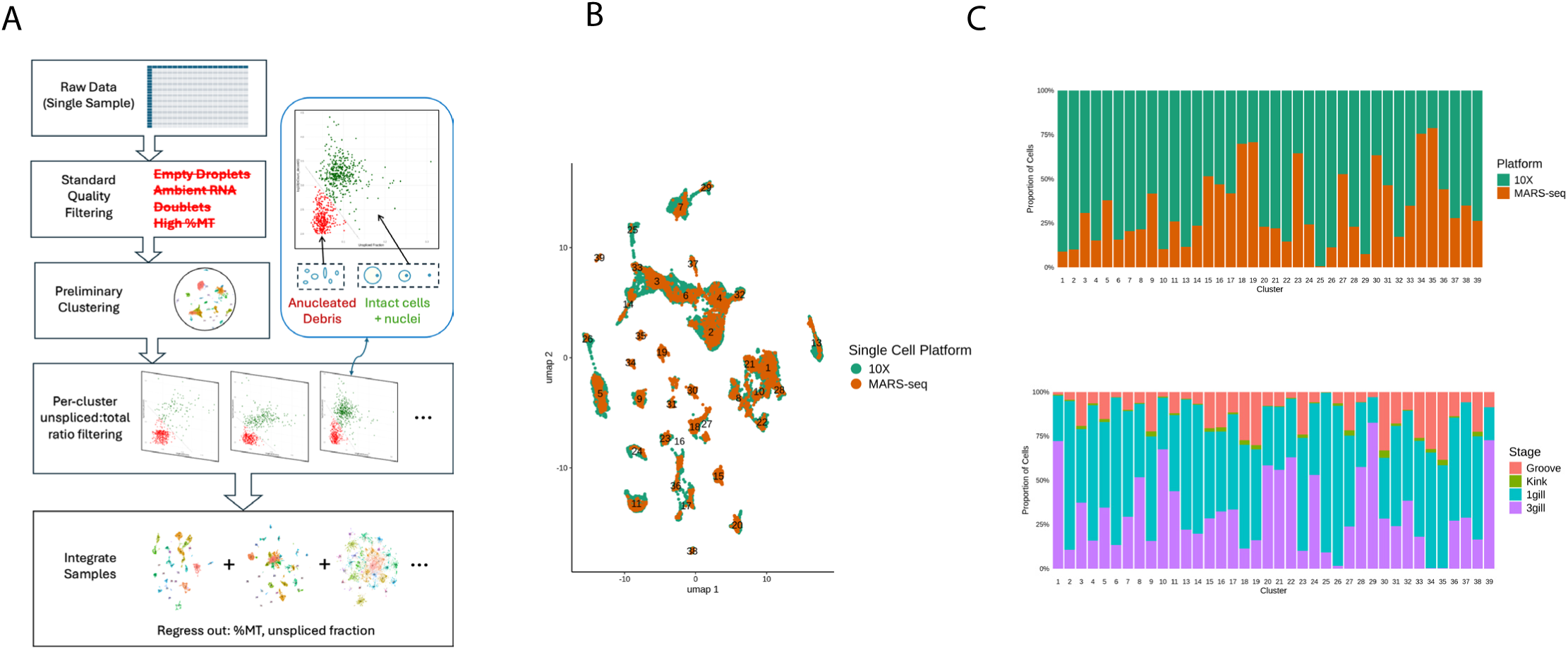
Processing and quality control of *Saccoglossus* whole body scRNA-seq data. A). Preprocessing pipeline used to filter and integrate whole body scRNA-seq data, including quality filtering steps based on fraction of UMIs that derive from unspliced transcripts. B). UMAP reduction plot showing single cell sequencing technology utilized to generate each cell. C). Proportion of cells in each cluster originating from each single cell sequencing technology (top) and each stage (bottom).

**Supplemental Figure 2.**
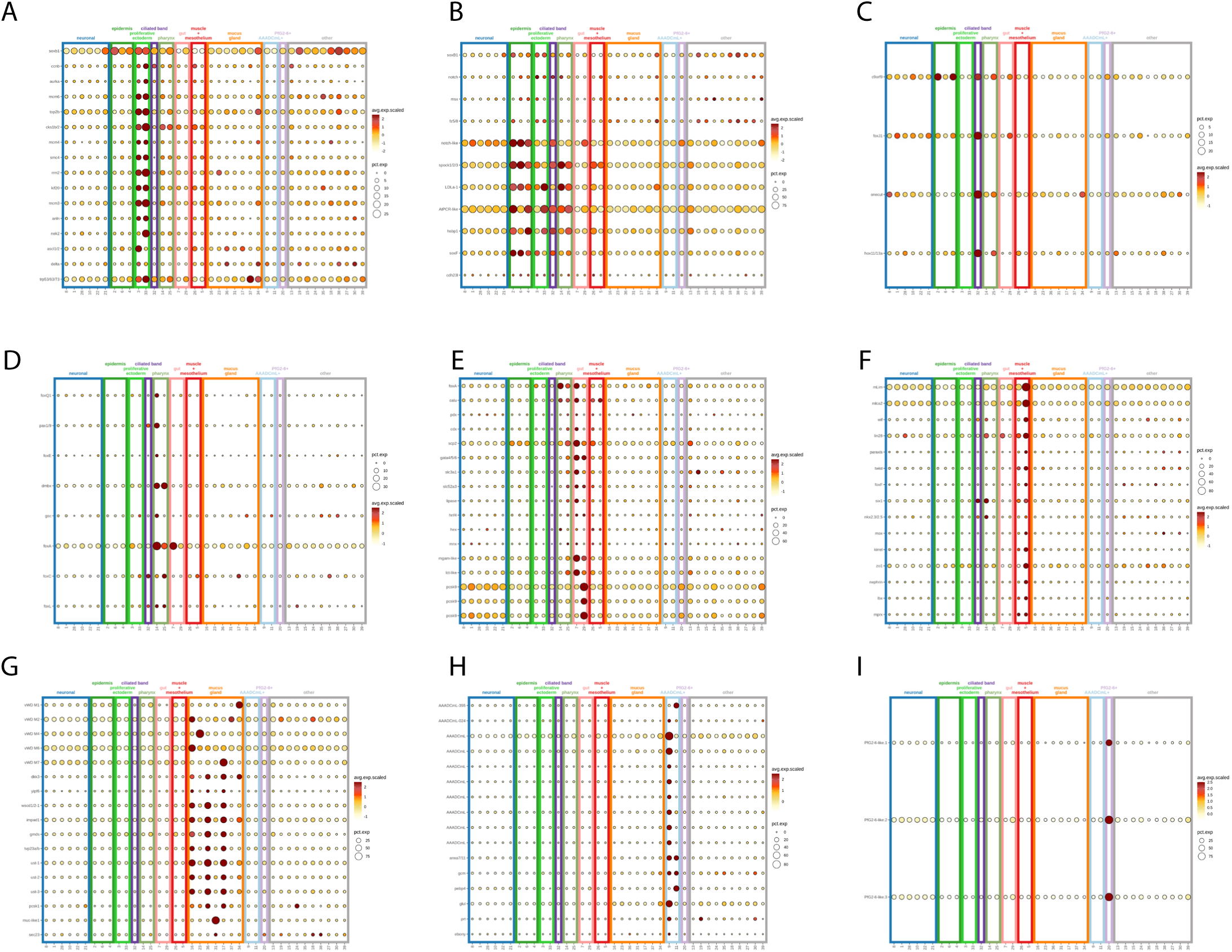
Cluster labeling of *Saccoglossus* whole body scRNA-seq data. A). Expression of markers used to identify proliferative ectoderm cells. B). Expression of markers used to identify epidermis cells. C). Expression of markers used to identify ciliated band cells. D). Expression of markers used to identify pharyngeal cells. E). Expression of markers used to identify gut cells. F). Expression of markers used to identify muscle + mesothelial cells. G). Expression of markers used to identify mucus gland cells. H). Expression of markers used to identify AAADCmL+ cells. I). Expression of markers used to identify PfG2-6+ cells.

**Supplemental Figure 3.**
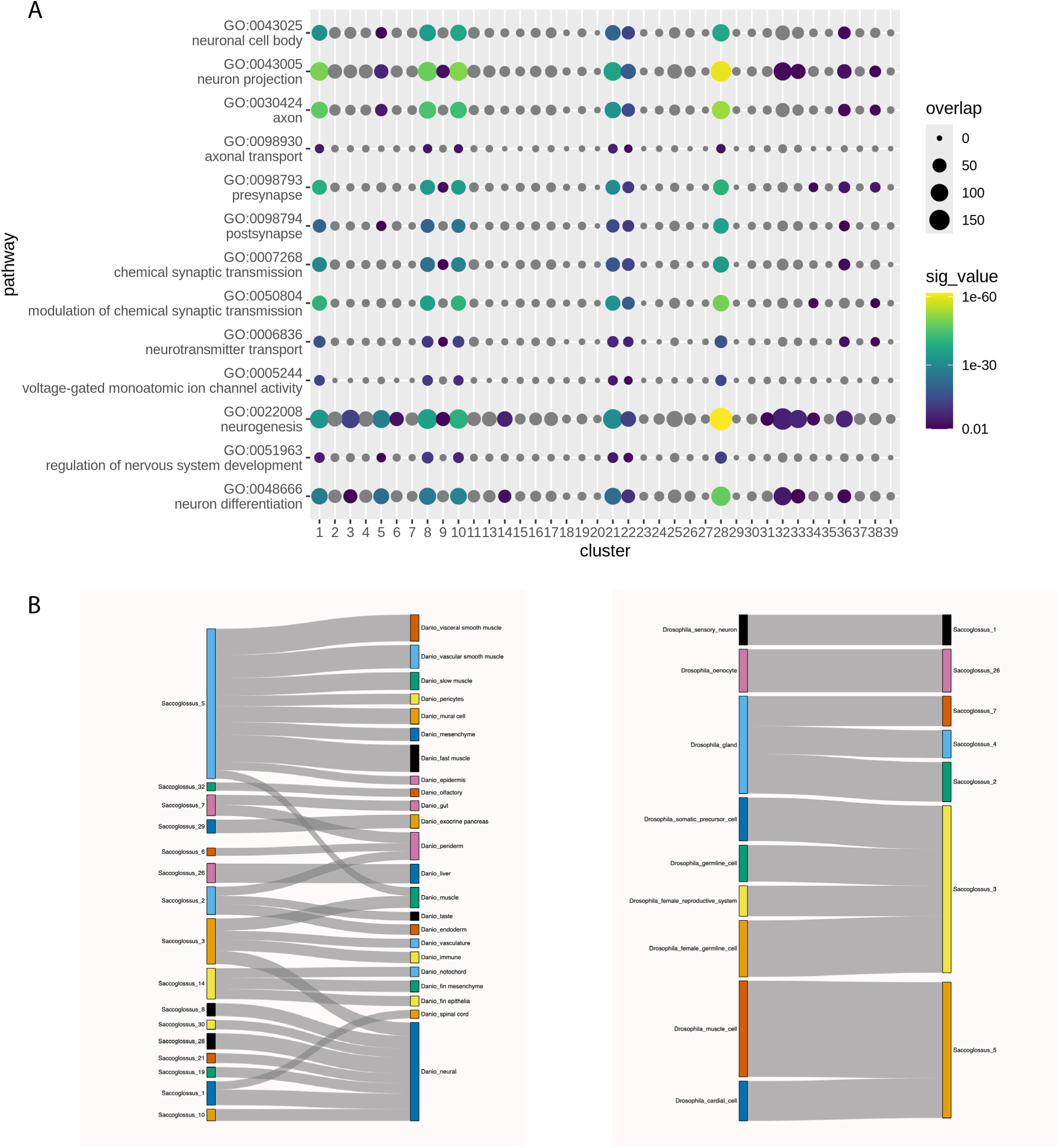
Orthogonal approaches for distinguishing neuronal and non-neuronal cells in the *Saccoglossus* whole body scRNA-seq dataset. A). Enrichment for neuron-associated GO terms amongst the marker genes for *Saccoglossus* whole body scRNA-seq clusters. B). Sankey plots summarizing results of SAMap comparison of gene expression in Saccoglosus whole body scRNA-seq clusters with data from the fly *Drosophila* (left) and the zebrafish *Danio* (right). Only associations supported by SAMap scores of above 0.2 are shown.

**Supplemental Figure 4.**
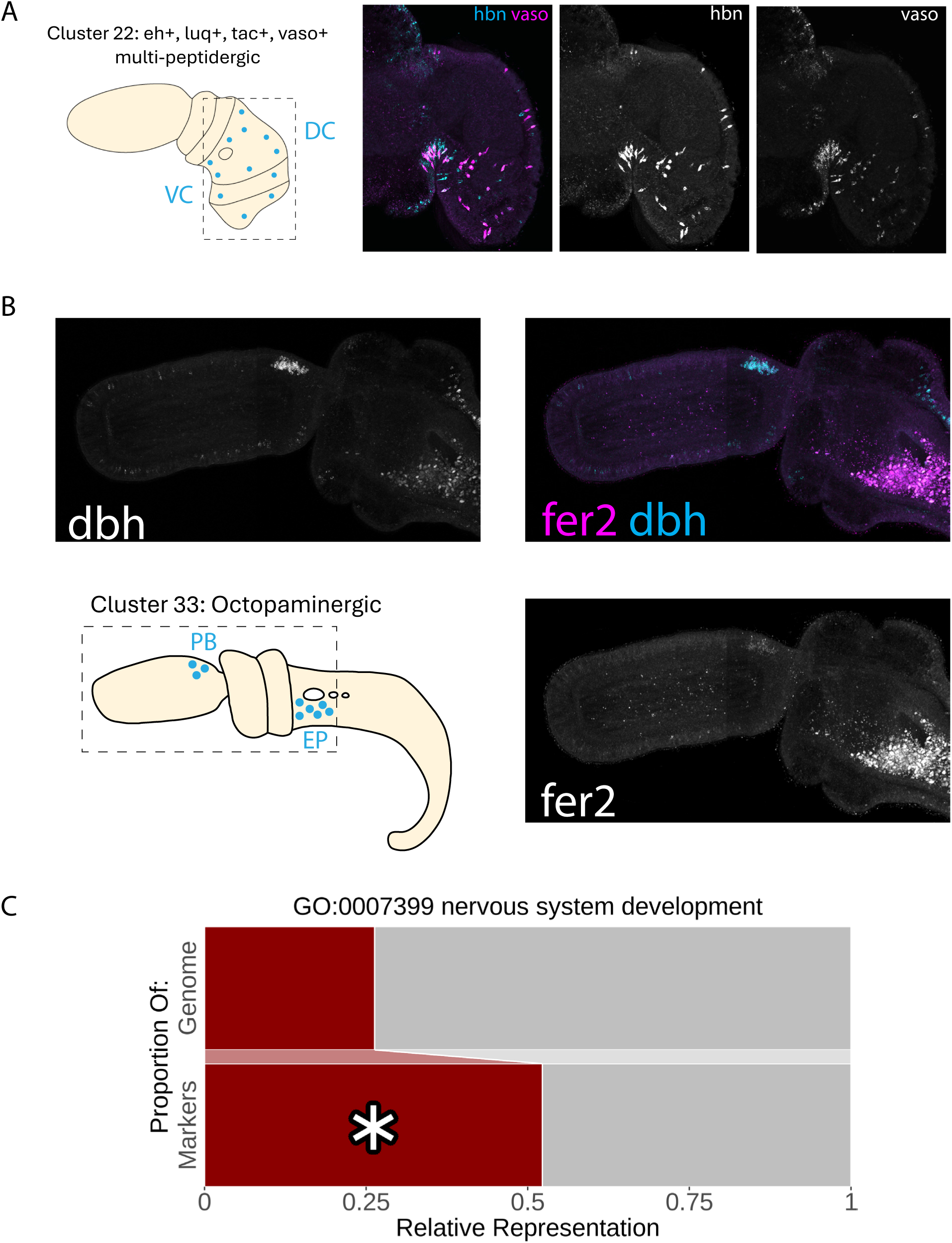
Additional characterization of the *Saccoglossus* neuronal TF regulatory code. A). Visualization of co-expression of TF *hbn* and effector *vaso*, associated with neuron cluster 22, *eh*+, *luq*+, *tac*+, *vaso*+ multi-peptidergic neurons. B). Visualization of expression of TF *fer2* in neuron cluster 33, octopaminergic neurons, as identified based on expression of effector *dbh*. C). Enrichment amongst neuronally-expressed TFs of those TFs associated with the GO term “nervous system development”.

**Supplemental Figure 5.**
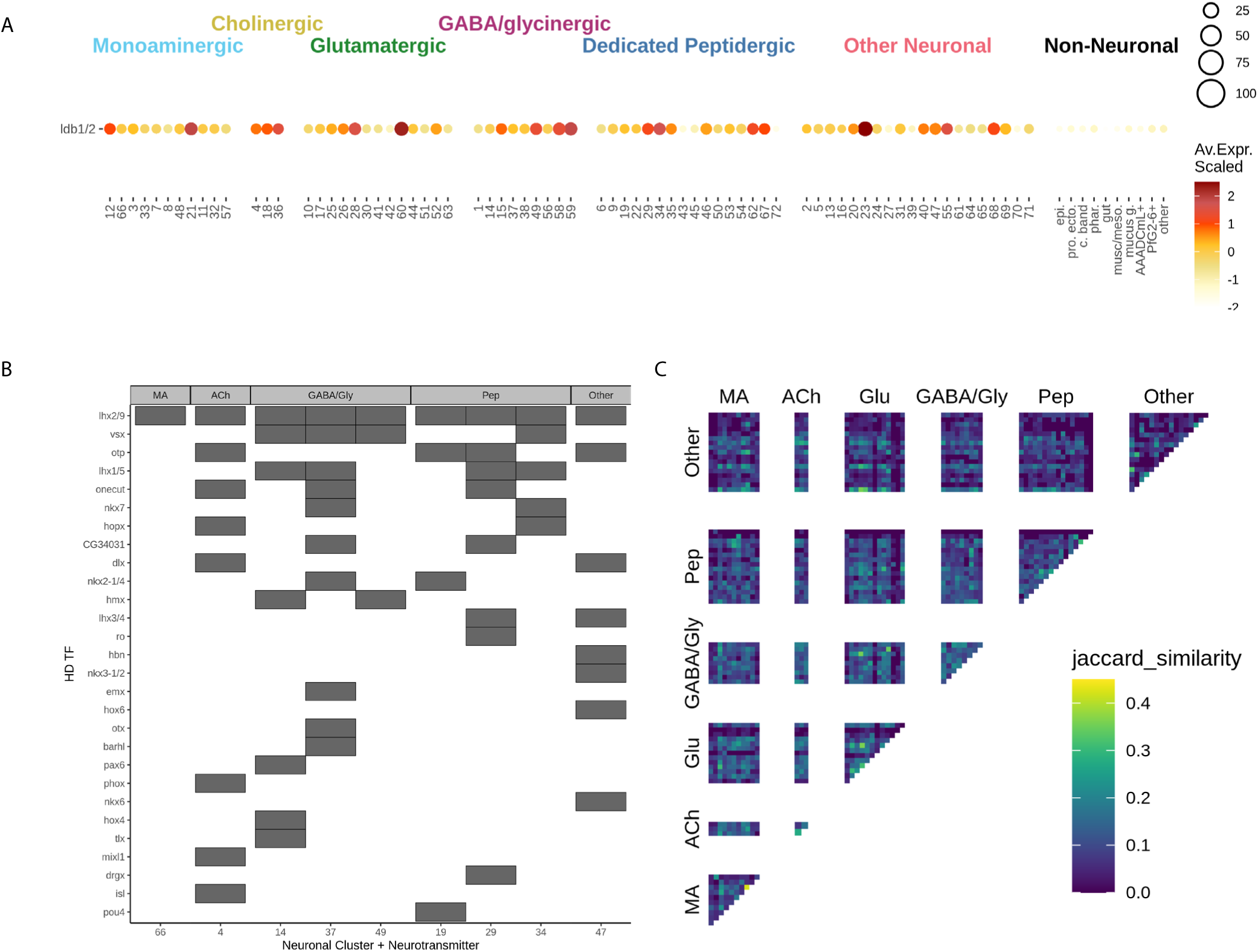
Additional characterization of the *Saccoglossus* neuronal TF regulatory code. A). Dotplot illustrating expression of LIM-HD cofactor *ldb1/2* in *Saccoglossus* neuronal subclusters and non-neuronal cell classes. B). Heatmaps demonstrating the combinations of HD TFs expressed in *lhx2/9*+ subclusters. C). Jaccard similarity of all *Saccoglossus* neuronal subclusters based on TF gene expression. Note that subclusters from the same neurotransmitter class do not tend to be more similar in their TF expression than subclusters expressing different neurotransmitters.

**Supplemental Figure 6.**
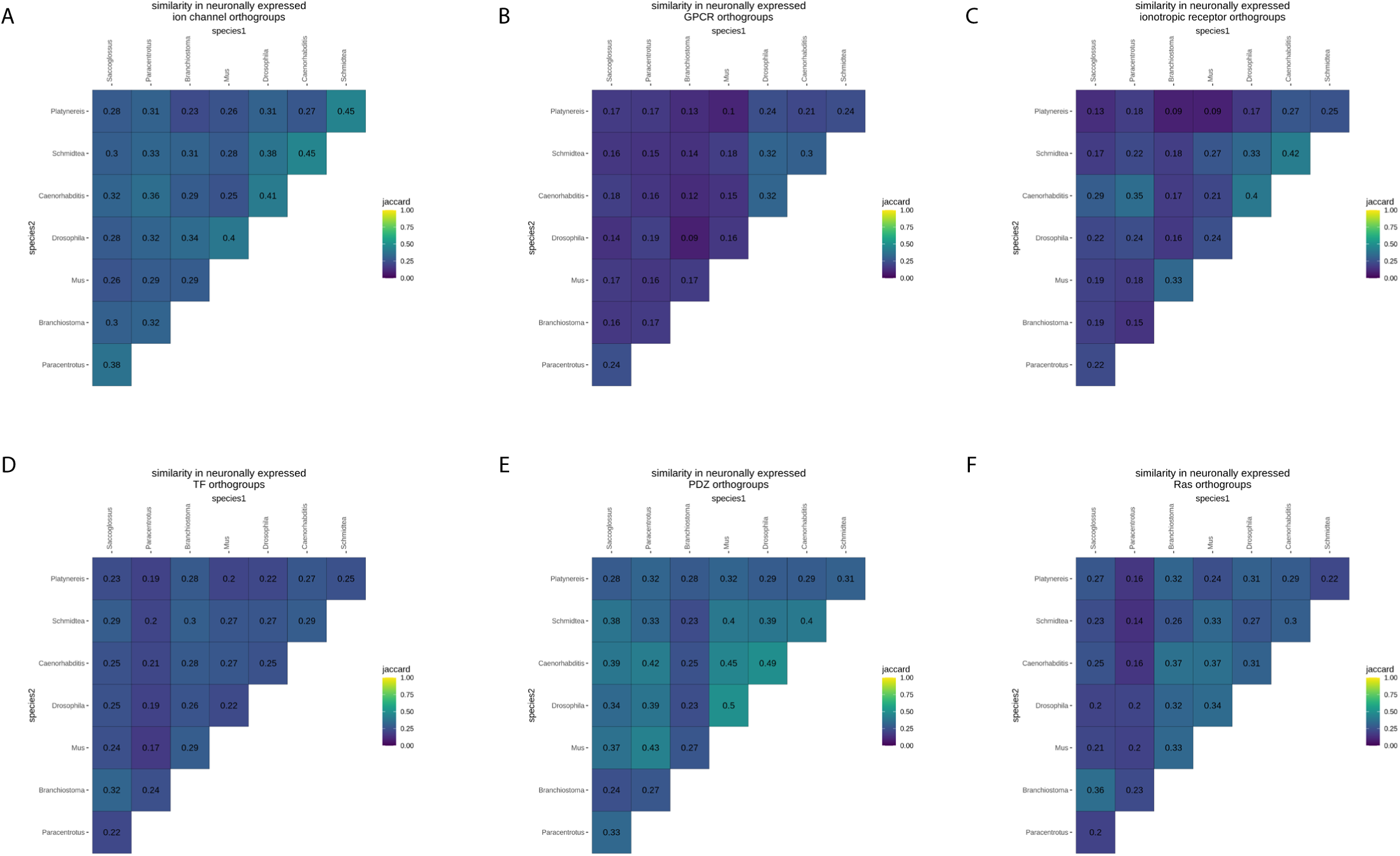
Jaccard similarity of neuronal gene sets across pairs of species, at the level of orthogroups. Orthogroup-level jaccard similarity for A). neuronally expressed ion channel genes, B). neuronally expressed GPCRs, C). neuronally expressed ionotropic receptors, D). neuronally expressed TFs, E). neuronally expressed PDZ-domain containing genes, and F). Ras genes.

**Supplemental Figure 7.**
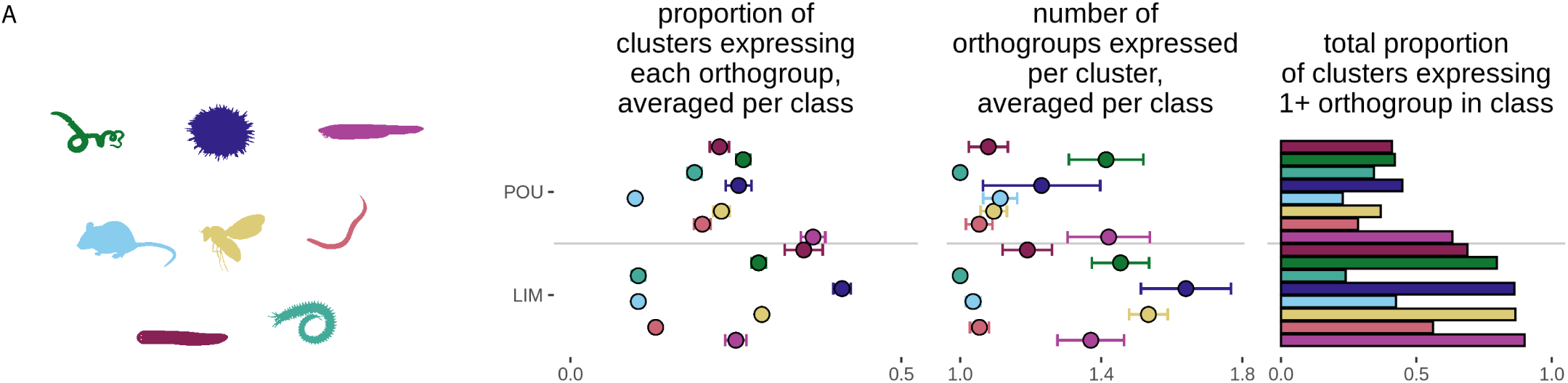
LIM- but not POU- HD TF genes show broad and complementary expression across multiple diverse bilaterians. A). Per-gene expression breadth, genes expressed per cluster, and combined expression breadth for LIM and POU HD TFs.

**Supplemental Figure 8.**
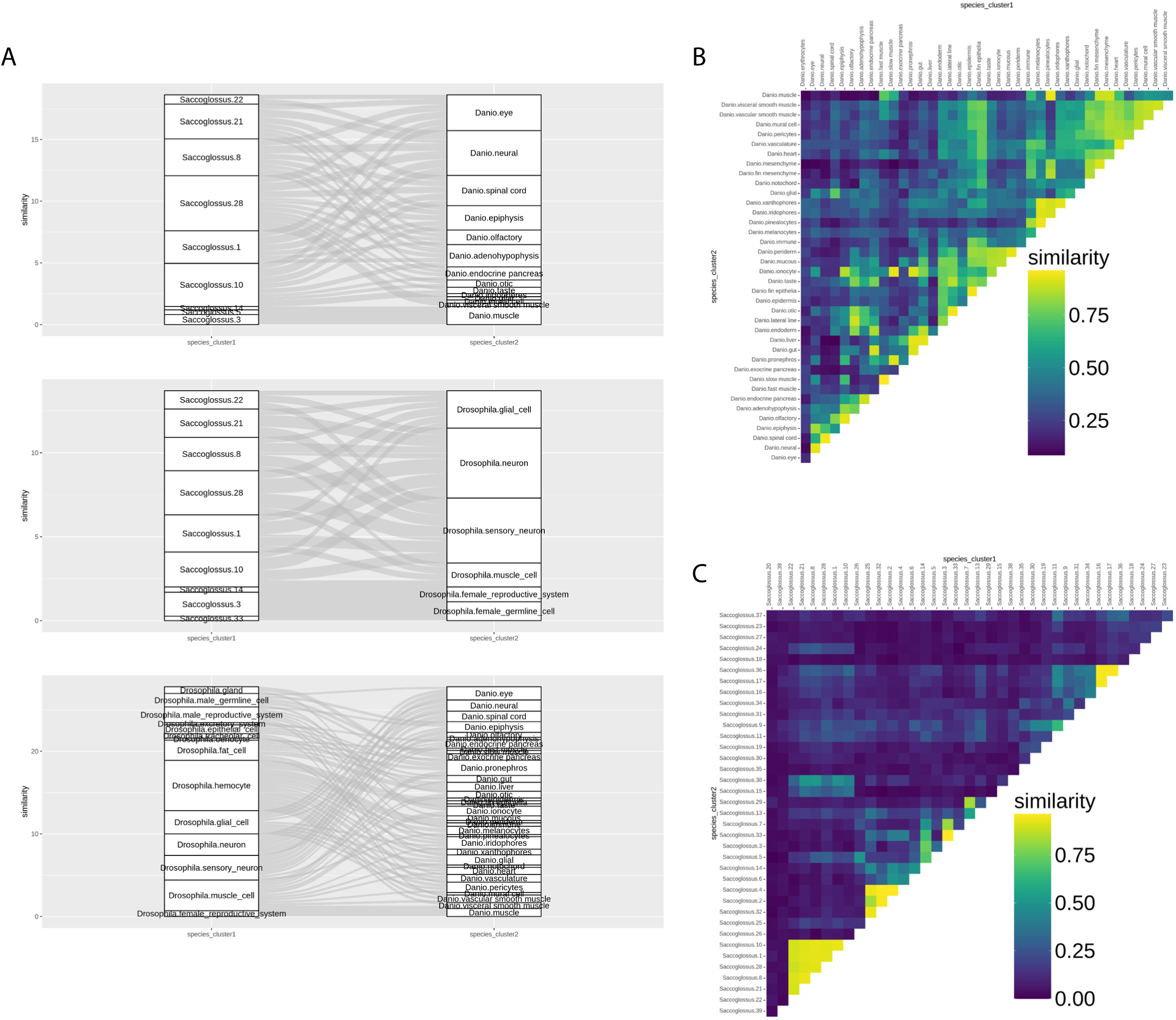
Validation of a specificity/set-based approach for quantifying similarities between clusters across and within species. A). Sankey plots summarizing results of specificity-weighted gene set score comparison of gene expression in whole body scRNA-seq clusters between data from *Saccoglossus* and *Danio* (top), *Saccoglossus* and *Drosophila* (middle) and *Danio* and *Drosophila* (bottom). Only associations supported by scores of above 0.25 are shown. B). Similarities between pairs of cell clusters within *Danio*. C). Similarities between pairs of cell clusters within *Saccoglossus*.

**Supplemental Figure 9.**
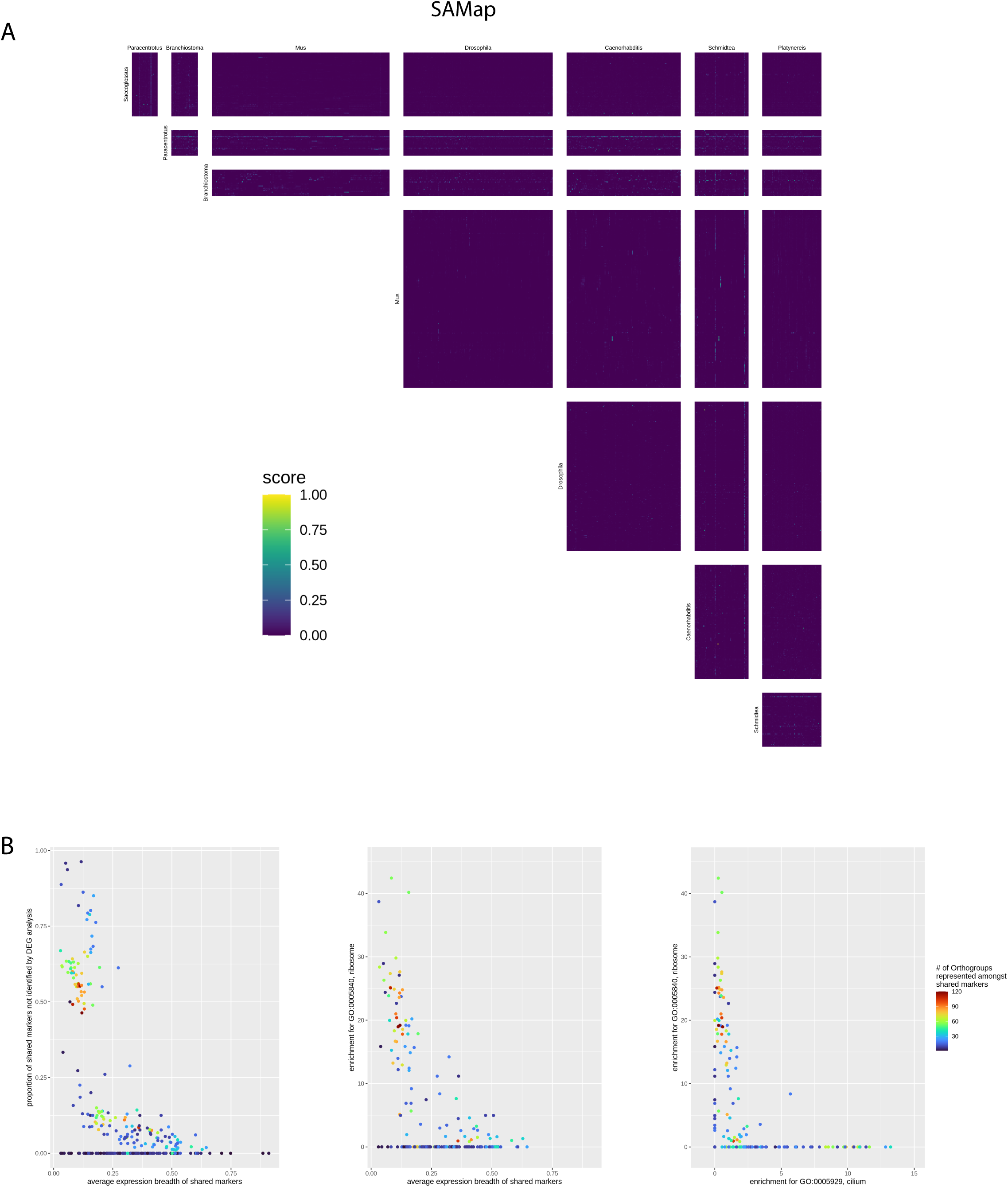
SAMap comparison of neuronal subclusters across all pairs of species. A). SAMap alignment scores. Note that most pairs of species demonstrate a small number of highly aligning clusters. B). For each SAMap high-scoring cross-species cluster alignment (score above), plots showing the relationships between number of shared markers identified, the breadth of expression of these markers, and number of these markers that are weak (not identifiable in our within-species differential expression analysis), enrichment for the "ribosome" GO term, and enrichment for the "cilium" GO term. Note that these confounding factors can account for the majority of alignments.

**Supplemental Figure 10.**
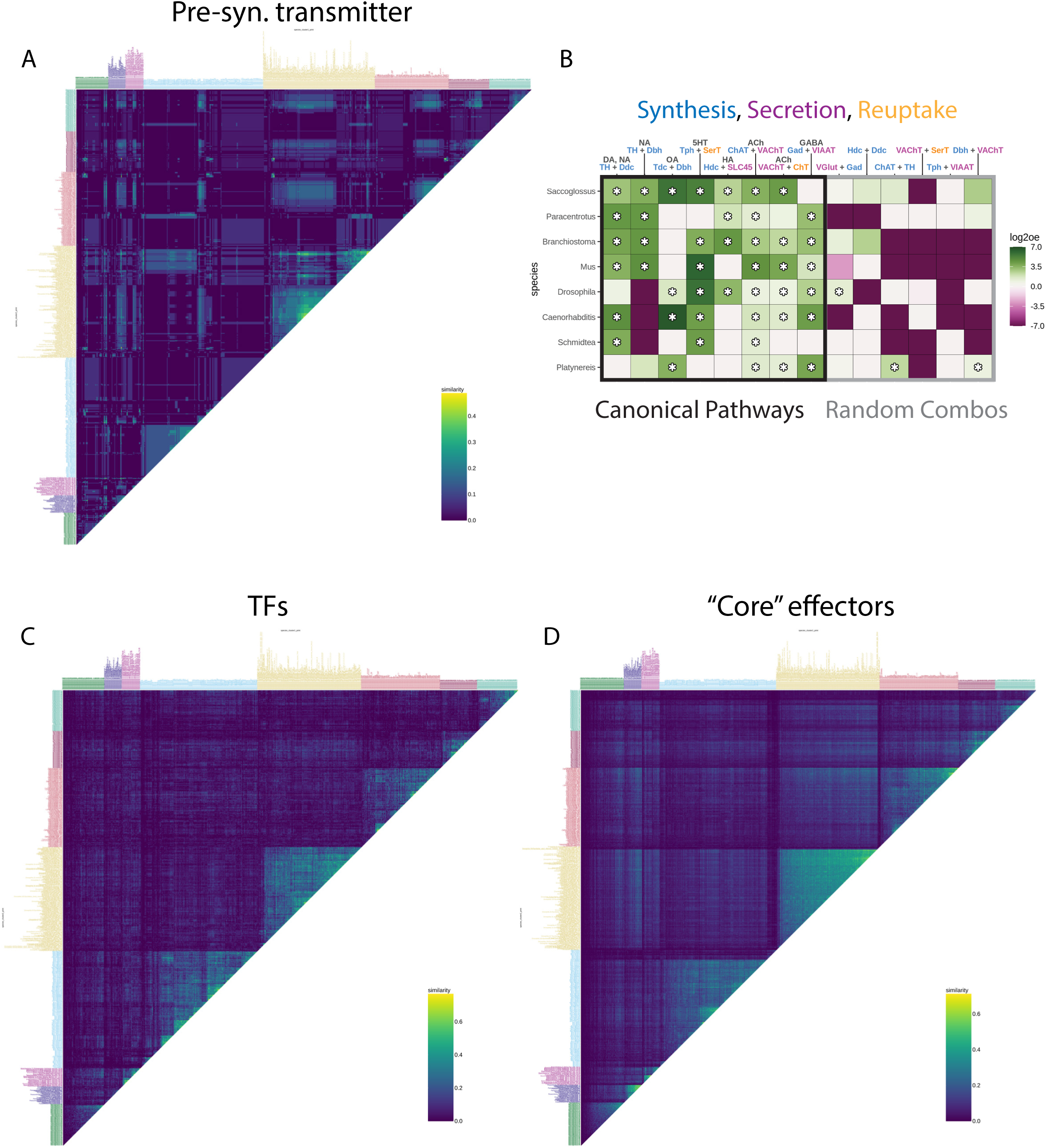
Specificity/set-based approach based quantification of neuronal subcluster similarity within and across species using only select sets of genes. A). Similarities when using only presynaptic small molecule neurotransmitter pathway genes. Note strong cross-species similarities, roughly corresponding to glutamatergic, GABAergic, cholinergic, and monoaminergic neurons. B). Cross-species comparisons of the patterns of co-expression amongst a few select example pairs of small molecule neurotransmitter pathway genes that are known to cooperate in the same pathway (left) vs. those that are not known to do so (right). The conserved patterns of co- and contra- expression drive the cross-species similarities observed in panel A. C). Similarities when using only TF genes. D). Similarities when using only “core” effectors (presynaptic neurotransmitter pathway genes, ion channels, ionotropic receptors, and GPCRs).

**Supplemental Figure 11.**
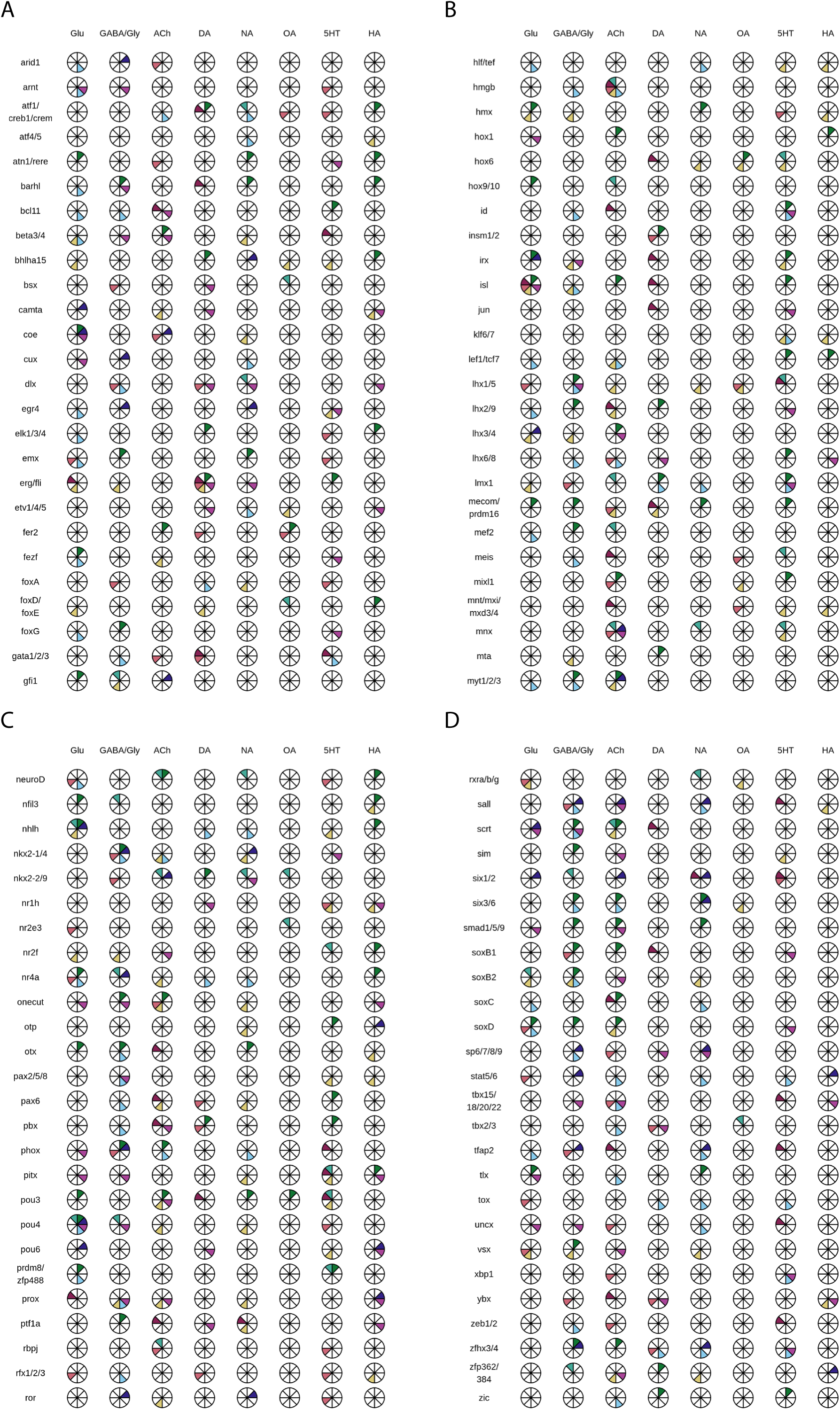
Associations between TFs and neurotransmitter identity associations. A-D). Neurotransmitter identity associations with reconstructed urbilaterian neuronal TFs as shown in Figure 4E. Color code is the same as in Figure 5D.

